# Data-driven system matrix manipulation enabling fast functional imaging and intra-image nonrigid motion correction in tomography

**DOI:** 10.1101/2024.01.07.574504

**Authors:** Peng Hu, Xin Tong, Li Lin, Lihong V. Wang

## Abstract

Tomographic imaging modalities are described by large system matrices. Sparse sampling and tissue motion degrade system matrix and image quality. Various existing techniques improve the image quality without correcting the system matrices. Here, we compress the system matrices to improve computational efficiency (e.g., 42 times) using singular value decomposition and fast Fourier transform. Enabled by the efficiency, we propose (1) fast sparsely sampling functional imaging by incorporating a densely sampled prior image into the system matrix, which maintains the critical linearity while mitigating artifacts and (2) intra-image nonrigid motion correction by incorporating the motion as subdomain translations into the system matrix and reconstructing the translations together with the image iteratively. We demonstrate the methods in 3D photoacoustic computed tomography with significantly improved image qualities and clarify their applicability to X-ray CT and MRI or other types of imperfections due to the similarities in system matrices.

## Introduction

Tomographic imaging modalities X-ray computed tomography (CT), magnetic resonance imaging (MRI), and photoacoustic computed tomography (PACT) produce cross-sectional images of tissue by detection of penetrating X-rays^1^, nuclear-magnetic-resonance-induced radio waves^2,3^, and light-absorption-induced ultrasonic waves^4^, respectively. Each modality with a certain setup is described by a system matrix^5–10^. Accurate image reconstruction poses requirements to the system matrix, which are often violated. For example, to achieve high temporal resolution for functional imaging, the spatial sampling density is often sacrificed, which introduces artifacts in the reconstructed image^11–13^ and may affect the functional signal extraction. Also, tissue motions such as heart beating^14–16^, breathing^17–19^, abdominal movement^20,21^, and fetal movement^22^, cause complex geometric errors in each system matrix, which introduce artifacts in the reconstructed image and compromise valuable image features.

Numerous methods have been proposed to compensate for system-matrix imperfections from image-domain^12,23–27^, signal-domain^28–33^, and cross-domain^34–38^ perspectives. However, due to the large size of each system matrix, these methods tend not to manipulate or correct the system matrix directly and have limitations. For sparse sampling functional imaging, traditional methods^12,25,26,32,38^ mitigate artifacts in images but their performances drop sharply as the sampling density reduces. Deep neural networks (DNNs)^27–29,31,34,37,39^ show high performance in mitigating artifacts but tend to generate false image features when the sampling density is low, and they require imaging-modality- and device-dependent datasets, which are not always available. Moreover, most of the methods introduce nonlinearity while mitigating artifacts, which disrupts the functional signals that are often much weaker than background signals.

For intra-image nonrigid motion correction, gating- and binning-based methods^16,30,33,35,36^ are commonly used. However, they require repeated data acquisition, which is time-consuming and infeasible for unrepeated motions. DNNs have also been used for motion correction^23,24^, however, they need specific training datasets that are not universally available, and it is challenging to reject falsely generated features in DNNs. Two system-matrix-level methods^40,41^ have been proposed for motion correction. In the first method^40,42^, the authors approximate general nonrigid motions with localized linear translations, identify possible motion paths from multichannel navigator data, and estimate the motion at each pixel using localized gradient-entropy metric in the image domain. However, quantifying localized motion from only navigator data is not robust, especially when the motion amplitude and noise level increase. In the second method^41^, the authors express breathing- and heartbeat-induced motions in basis functions by performing singular value decomposition (SVD) and resolve these motions in imaging. However, for general motions, especially unrepeated motions, the method’s performance is unknown. The high computation cost of SVD in the method also restricts its application to 3D imaging.

Here, we compress the system matrices using SVD and fast Fourier transform (FFT), which enables efficient system matrix slicing and manipulation. Then, we use two new methods for functional imaging and motion correction, respectively, by performing data-driven manipulations of these matrices. Both methods are applicable to CT, MRI, and PACT. For sparse sampling functional imaging, we incorporate a prior image into the system matrix to reduce unknown variables in image reconstruction. Special configurations in the method maintain linearity in image reconstruction while substantially mitigating artifacts, which is critical for weak functional signal extraction. For intra-image nonrigid motion correction, we approximate the motion with localized linear translations^40^. Starting from an initially reconstructed motion-blurred image, we first estimate the translation of each subdomain of the object by minimizing the difference between the simulated signals from the subdomain and the detected signals. With the estimated translations, we update the system matrix and reconstruct the image again. We iterate the correction-reconstruction process to obtain the final image. This method does not require repeated data acquisition and is effective for unrepeated motions.

In this work, we use 3D PACT for demonstration due to its representatively large size in tomography: light-absorption-induced ultrasonic wave from every voxel in an image is detected by every transducer element and the system matrix is intrinsically a 6D tensor^10^. We apply the proposed methods to both numerical simulations and *in vivo* experiments: mouse brain functional imaging with sparse sampling and intra-image motion correction in human breast imaging. We demonstrate that both methods substantially improve the functional and structural image qualities.

## Results

### System matrix compression based on SVD and FFT

In this study, we use the 3D PACT system reported previously by Lin et al.^43^, which consists of four 256-element arc transducer arrays (central frequency of 2.25 MHz and one-way bandwidth of 98%), and we assume a homogenous medium. The rotation of the four arc-arrays forms a virtual hemispherical array truncated at the bottom for light delivery (marked as blue arcs in **Fig. 1a**), and the detected ultrasonic signals are used to form a 3D image (marked as a rectangular cuboid). We denote the number of voxels in the image and the number of virtual elements in the 2D array as *M* and *N*, respectively. The system matrix is formed by *MN* responses of all virtual elements to the signals from all voxels, with three responses’ paths shown as black-dotted arrows in **Fig. 1a**. It is expensive to store and manipulate these responses directly from memory and computation perspectives, respectively. Considering that the responses’ arrival times are determined by the voxel-element distances, we temporally shift the responses so that the arrival times are aligned along the blue-dotted line in **Fig. 1b**. Performing grouped SVDs to the shifted responses, we approximate these responses as linear combinations (coefficients shown in **Fig. 1b**) of three temporal singular functions with high accuracy: an efficient compression of the system matrix. In **Supplementary Note 1**, we elaborate on the process of performing SVD to the responses grouped for each virtual element in the element’s local coordinate system. Due to the homogeneous-medium assumption and identity of all elements in the system, SVDs for all virtual elements share the same set of temporal singular functions, with the first three shown as red, gray, and blue curves, respectively, in **Fig. 1b**. In **Supplementary Fig. 1e–i**, we demonstrate that the arrival-time-dependent temporal shifting is essential for the compression based on SVD.

**Fig. 1.**
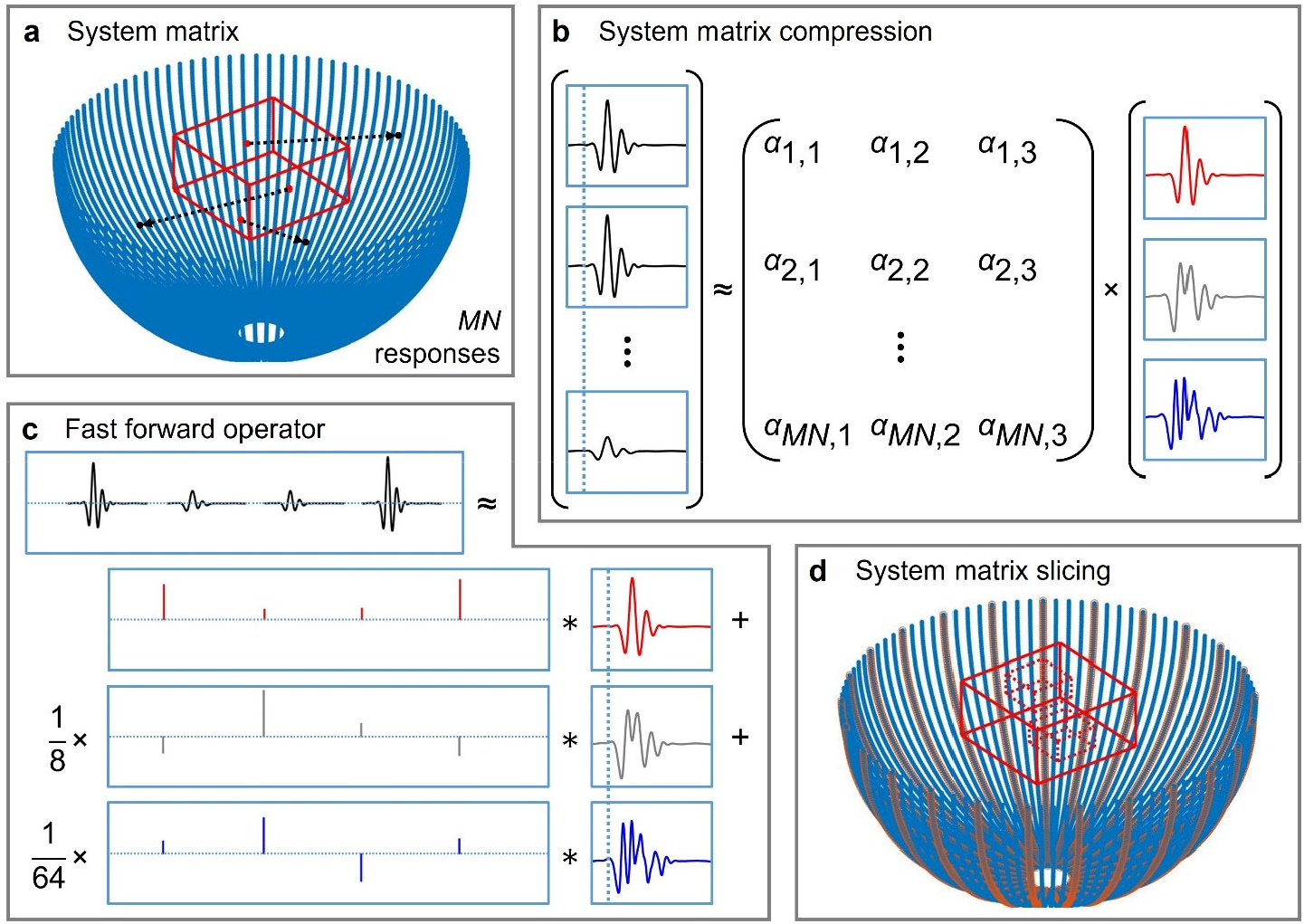
System matrix compression and slicing. **a** A virtual 2D array (blue curves) formed by rotation of four arc arrays, the image domain (a rectangular cuboid with red-solid edges), and paths of three responses marked by black-dotted arrows from voxels (red dots) to transducer elements (black dots). **b** Responses temporally shifted to remove the differences in arrival times (aligned along the blue-dotted line) from the voxels to elements, and compression of these responses based on SVD with the first three temporal singular functions shown as red, gray, and blue curves, respectively. **c** Signals detected by an element approximated by three convolutions with the three temporal singular functions (zero time indicted by the blue-dotted line), respectively. **d** System matrix slicing in voxel indices (rectangular cuboids with red-dotted edges) and virtual element indices (blue arcs with red boundaries).

The system matrix compression not only saves memory in storing the matrix but also enables fast and accurate forward simulation. For each combination of a voxel and a virtual element, instead of adding the whole weighted response to the signals, we only access three coefficients: values of the first three spatial singular functions. After obtaining all the coefficients with different arrival times for each virtual element, we perform three convolutions (implemented by FFT) to obtain the signals, as shown in **Fig. 1c**. We summarize the implementations of the original and compressed system matrices in Eqs. (1) and (2), respectively. Through forward simulations, we demonstrate that the compressed system matrix is approximately 42 times as fast as the original system matrix with negligible signal-domain relative errors (maximum value of 0.005) for this study (**Supplementary Fig. 2**). Further, we reconstruct images using a nonnegativity-constrained iterative method (Eq. (S33)) with the compressed system matrix from signals simulated using the original system matrix, and observe negligible image-domain relative errors (maximum value of 0.007) for this study (**Supplementary Fig. 3**).

The implementation of the system matrix based on Eq. (2) is not only efficient but also explicit, meaning subsets of the system matrix are directly accessible. For example, to obtain the signals from *M*′ voxels (enclosed in two rectangular cuboids with red-dotted edges in **Fig. 1d**) detected by *N*′ virtual elements (blue arcs with red boundaries in **Fig. 1d**), we can access the relevant system matrix elements directly and finish the computation in 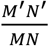 of the time for *M* voxels and *N* elements. It needs to be noted that this scaling property critical for this study is invalid for wave-equation-based implementations^9^ of the system matrix.

### Sparse sampling functional imaging using a hybrid method

We propose a hybrid method for functional imaging (Methods, **Supplementary Fig. 6**) by combining an iterative method with the universal back-projection (UBP) method^44^. We first demonstrate the performance of the hybrid method by using it to reconstruct images from signals of a numerical phantom acquired by virtual arrays with different numbers of arcs: 4*N*_loc_ = 76, 40, 28, 20, 16, 12. Here *N*_loc_ is the number of rotating locations of the four-arc array (with 128 transducer elements in each arc) in a virtual array. Images of the numerical phantom reconstructed using UBP, the regularized iterative method (Eq. (S34)), and the hybrid method (Eq. (5) with a prior image obtained by performing a smooth modulation to the numerical phantom) are shown in the first three columns in **Fig. 2a**. The used virtual arrays are shown as blue arcs with red boundaries in the fourth column in **Fig. 2a**. We discuss the selection of regularization parameter in **Supplementary Note 6** and, specifically, the best regularization parameters for different values of 4*N*_loc_ (**Supplementary Fig. 5**) that are used in the iterative method in **Fig. 2a**. Maximum amplitude projections (MAPs) of these images along the *z*-axis are shown in **Supplementary Fig. 7**. We see that, as 4*N*_loc_ decreases from 76 to 12, the artifacts in the images reconstructed using UBP become more abundant. The regularized iterative method mitigates the relatively weak artifacts in all images but failed to suppress the strong artifacts such as in the images with 4*N*_loc_ = 16, 12. In contrast, with the help of the prior image, the hybrid method significantly mitigates the artifacts. Quantitatively, for each method, we calculate the structural similarity index measures (SSIMs) between the images with 4*N*_loc_ = 40, 28, 20, 16, 12 and that with 4*N*_loc_ = 76, and compare the values in **Fig. 2b**. The hybrid method performs the best in mitigating artifacts and maintaining true features.

**Fig. 2.**
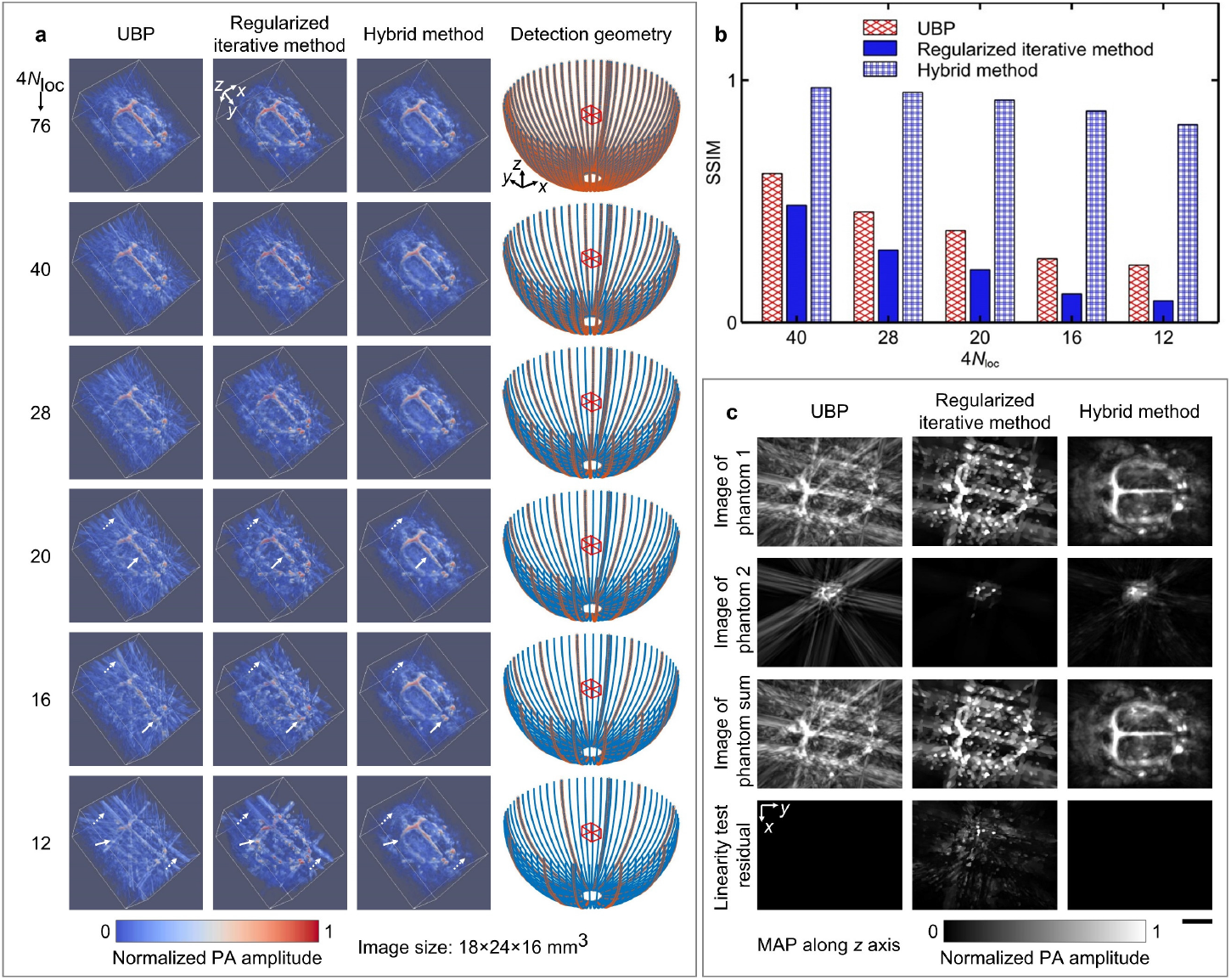
UBP, regularized iterative method, and the proposed hybrid method for sparse-sampling imaging, and their linearity tests. **a** Images reconstructed by the three methods (first three columns) from signals detected at sparsely distributed elements (red-bounded blue curves in the fourth column) for 4*N*_loc_ = 76, 40, 28, 20, 16, 12 . Examples of maintained features and suppressed artifacts are indicated by white-solid and white-dotted arrows, respectively. **b** SSIMs between the reconstructed images with 4*N*_loc_ = 40, 28, 20, 16, 12 and those with 4*N*_loc_ = 76 for the three methods. **c** Linearity tests of the three methods for 4*N*_loc_ = 12. Scale bar, 5 mm.

Additionally, we test the linearity of each method by reconstructing two numerical phantoms and their summation (shown in **Supplementary Fig. 8a–c**, respectively). MAPs of the reconstructed images and the linearity test residuals (image of the summation subtracted by images of the two phantoms) of the three methods with 4*N*_loc_ = 76, 12 are shown in **Supplementary Fig. 8e-f** and **Supplementary Fig. 9c**, respectively, which validate that the hybrid method maintains linearity while mitigating artifacts. We summarize the MAPs in the linearity tests with 4*N*_loc_ = 12 in **Fig. 2c**. In another test of the hybrid method using a prior image with dot artifacts (**Supplementary Fig. 9b**), the linearity is still valid although artifacts appear in the reconstructed images (**Supplementary Fig. 9d**). In summary, UBP and the hybrid method are linear, but the regularized iterative method is nonlinear. In the hybrid method, the prior image quality does not affect the linearity but affects the reconstructed image quality.

Due to the requirement of a prior image, the hybrid method’s practical value mainly lies in fast functional imaging with sparse sampling. We perform numerical simulations of functional imaging with 4*N*_loc_ = 12 in **Supplementary Note 7**. After reconstructing images from simulated signals using the three methods, we extract functional signals from them using a method based on regularized correlation (Eq. (7)) and form functional images. As shown in **Supplementary Fig. 11** and **Supplementary Video 1**, artifacts in the UBP-reconstructed images cause artifacts in the functional images, the regularized iterative method mitigates artifacts in functional images but also compromises the true functional region, and the proposed hybrid has the best performance.

Moreover, we apply UBP, the regularized iterative method, and the hybrid method to mouse brain functional imaging *in vivo* using the four-arc system. We first obtain a prior image of a mouse brain through dense sampling (4*N*_loc_ = 396), then electrically stimulate its right front paw and continuously acquire signals from the mouse brain through sparse sampling (4*N*_loc_ = 76, 2 s per image). We use subsets of the sparsely sampled signals (4*N*_loc_ = 40, 20, 12) to demonstrate the performance of the hybrid method. For one set of sparsely sampled signals, the images reconstructed using UBP, the regularized iterative method, and the hybrid method for 4*N*_loc_ = 40, 20, 12 are shown in **Fig. 3a**. We observe that the iterative method mitigates the artifacts (e.g., those indicated by white-dotted arrows) but compromises low-amplitude features (e.g., those indicated by white-solid arrows for 4*N*_loc_ = 20). In contrast, the hybrid method maintains low-amplitude features while substantially mitigating the artifacts, resulting in images more similar to the densely sampled image. Electrical stimulation of the mouse’s right front paw occurs in five cycles, each with 12-s stimulation on and 12-s off, as shown in **Fig. 3b**. To find the best regularization parameter (*λ*_f_) in the regularized correlation (Eq. (7)), we obtain functional images from the images reconstructed through the three methods for 4*N*_loc_ = 40, 20, 12 using *λ*_f_ = 0.02, 0.08, 0.32, 1.28, 5.12. For UBP and the hybrid method, *λ*_f_ = 0.32 is the best choice to maintain the true functional region and suppress false positive regions, and the functional images for 4*N*_loc_ = 12 are shown in **Supplementary Fig. 12**. For the regularized iterative method, we compare the images for 4*N*_loc_ = 40, 20, 12 in **Supplementary Fig. 13** and observe that *λ*_f_ = 0.08 is the best choice. We summarize the obtained functional images with best values of *λ*_f_ in **Fig. 3c** and **Supplementary Video 2**. Results from UBP and the hybrid method match well for 4*N*_loc_ = 40. The hybrid method is slightly (significantly) better than UBP for 4*N*_loc_ = 20 (4*N*_loc_ = 12). Due to the violation of linearity, the regularized iterative method compromises the true functional region: leading to its shrinkage for 4*N*_loc_ = 40, 20 and its decimation altogether for 4*N*_loc_ = 12. In summary, the proposed hybrid method enables fast functional imaging with highly sparse sampling.

**Fig. 3.**
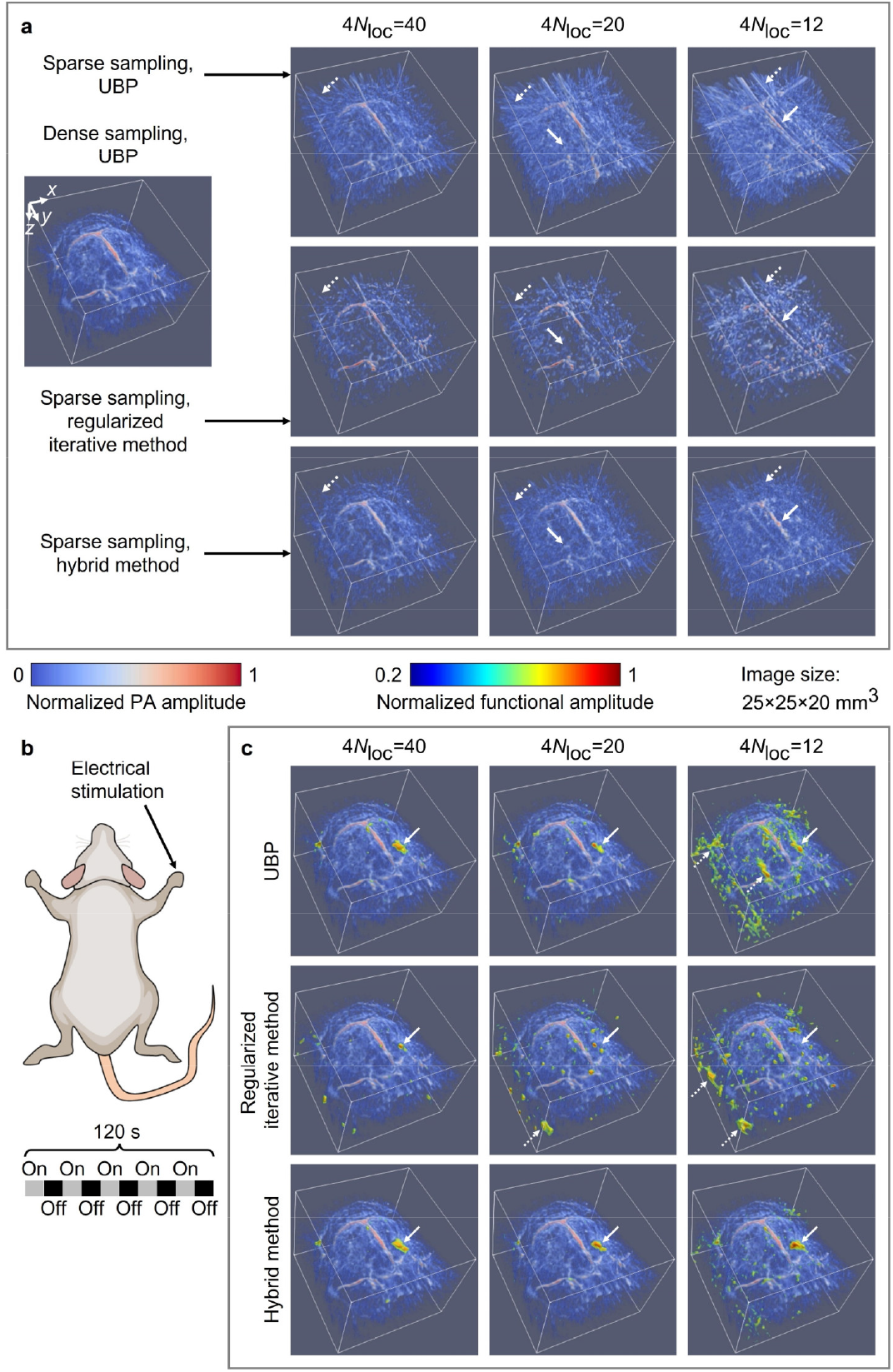
Sparse-sampling mouse brain functional imaging *in vivo*. **a** A densely sampled image of a mouse brain reconstructed by UBP (left column, 4*N*_loc_ = 396) and sparsely sampled images of the mouse brain reconstructed using UBP (first row), the regularized iterative method (second row), and the hybrid method (third row), respectively, for 4*N*_loc_ = 40, 20, 12 . Examples of suppressed artifacts and maintained features are indicated by white-dotted and white-solid arrows, respectively. **b** Electrical stimulation to the mouse’s right front paw: five cycles, each with 12-s stimulation on and 12-s off. **c** Functional images obtained from the images reconstructed using UBP (first row, *λ*_f_ = 0.32), the regularized iterative method (second row, *λ*_f_ = 0.08), and the hybrid method (third row, *λ*_f_ = 0.32), respectively, for 4*N*_loc_ = 40, 20, 12. The true functional regions in all images are indicated by white-solid arrows, and examples of false positive regions are indicated by white-dotted arrows.

### Intra-image nonrigid motion correction

We propose a method for intra-image nonrigid motion correction through data-driven manipulation of the system matrix (Methods, **Supplementary Note 8**), with the workflow shown in **Supplementary Fig. 14**. We first demonstrate the method in numerical simulations. From a numerical phantom (**Fig. 4a**) obtained in 3D imaging of a human breast, we simulate numerical phantoms with translation- and deformation-induced intra-image motions, whose patterns are defined by Eqs. (9) and (10), respectively. We depict the array rotation with tissue translation and deformation in **Fig. 4b-c**, respectively. A 90° rotation of the four-arc array (*N*_loc_ = 72, *N*_ele_ = 4 × 72) allows the transducer elements to detect signals needed to form a 3D image, during which tissue motions occur. The amplitudes of translation and deformation are controlled by *A*_tra_ and *A*_def_, respectively, and the number of periods of the motion during a 90° rotation of the array is denoted as *N*_per_. We visualize in **Supplementary Video 3** the synchronized motions of the array and numerical phantom using two examples: *A*_tra_ = 1.2 mm, *N*_per_ = 3 and *A*_def_ = 0.12, *N*_per_ = 3 for translation and deformation, respectively. For detected signals in this video, we reorganize the *N*_loc_ locations of the four-arc array (*N*_ele_ elements) to 4*N*_loc_ locations of the single arc array (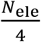 elements) with new indices 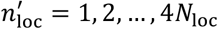 and 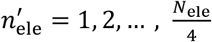. The signals for 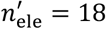 and 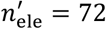 (closer to the array-rotation axis) are visualized in the video.

**Fig. 4.**
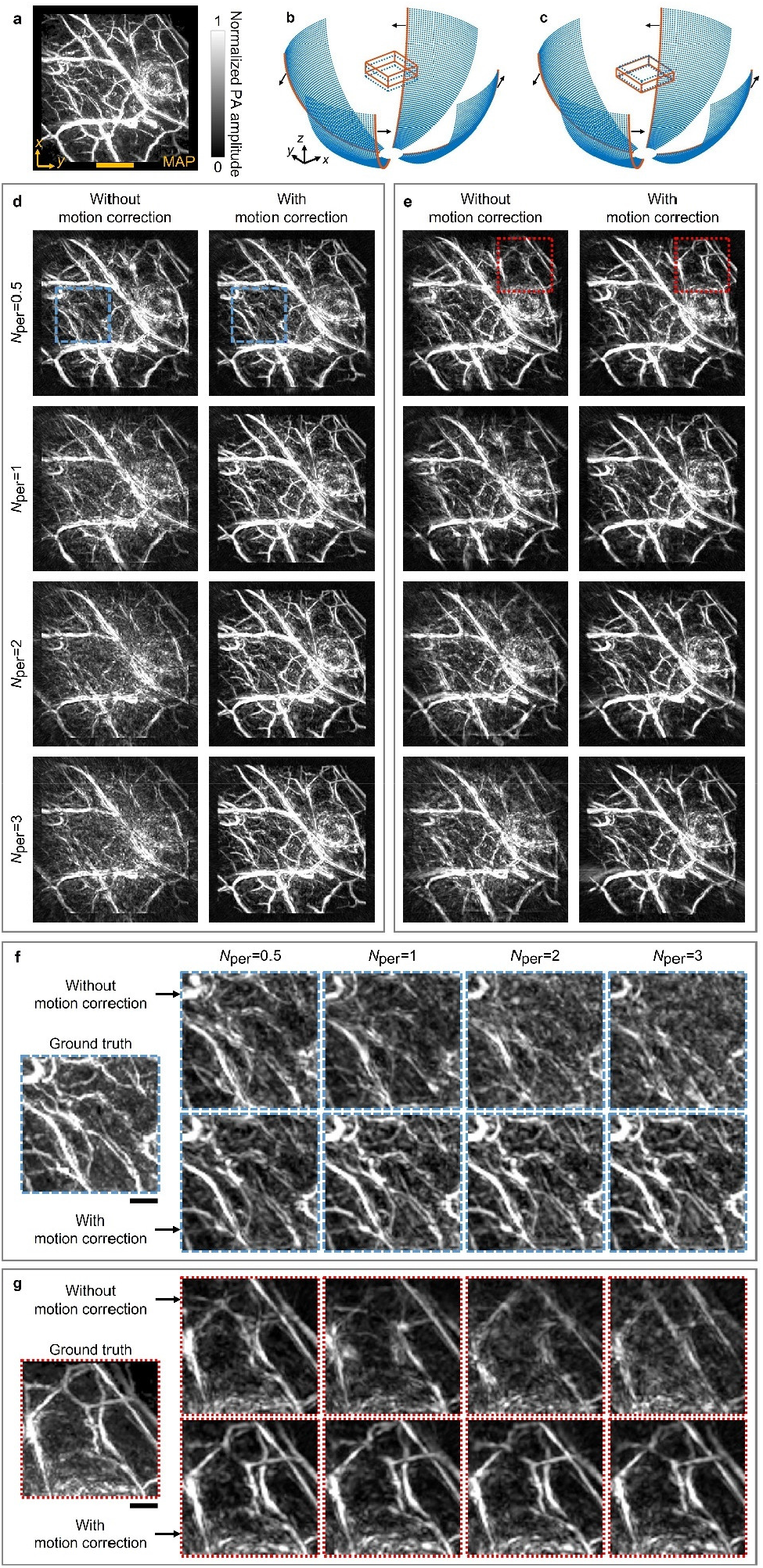
Intra-image nonrigid correction of motions induced by tissue translation and deformation. **a** MAP of an image (a human breast image subset with a volume of 44.8×44.8×14.4 mm^3^) along the *z*-axis. Scale bar, 1 cm. **b-c** Depictions of the intra-image motions induced by tissue translation and deformation, respectively. The image translates (deforms) from a rectangular cuboid with blue-dotted edges to one with red-solid edges as the four-arc array rotates, with the passed virtual element locations marked as blue dots, the latest element locations marked as red curves, and the rotation direction indicated by black arrows. **d-e** MAPs of the images reconstructed from the signals with motions induced by tissue translation (*A*_tra_ = 0.6 mm) and deformation (*A*_def_ = 0.06), respectively, without and with motion correction for *N*_per_ = 0.5,1,2,3 . **f-g** Closed-up subsets of the MAPs enclosed in blue-dashed boxes (in **d** and **f)** and red-doted boxes (in **e** and **g**), respectively, with the ground truths.

Additionally, we visualize the translation-induced motions with detected signals for *A*_tra_ = 1.2 mm, *N*_per_ = 0.5, 1, 2, 3 in **Supplementary Video 4**, and the deformation-induced motions with detected signals for *A*_def_ = 0.12, *N*_per_ = 0.5, 1, 2, 3 in **Supplementary Video 5**.

We reconstruct images from the simulated signals without and with motion correction and compare their MAPs along the z-axis in **Supplementary Video 6** (translation-induced motions with *N*_per_ = 0.5, 1, 2, 3 and *A*_tra_ = 0.2, 0.4, 0.6, 0.8, 1.0, 1.2 (mm)) and **Supplementary Video 7** (deformation-induced motions with *N*_per_ = 0.5, 1, 2, 3 and *A*_def_ = 0.02, 0.04, 0.06, 0.08, 0.10, 0.12). The proposed motion correction method improves image quality for every set of signals. We show those MAPs for *N*_per_ = 0.5, 1, 2, 3 in **Fig. 4d** (translation, *A*_tra_ = 0.6 mm) and **Fig. 4e** (deformation, *A*_def_ = 0.06). Moreover, we pick subsets of the MAPs in blue-dashed box and red-dotted box in **Fig. 4d** and **e**, respectively, and compare closeups of them in **Fig. 4f** and **g** with ground-truth MAPs. We also compare those closeups for *N*_per_ = 0.5, 3 in **Supplementary Video 8**. We see that, motion correction not only reduces the motion-induced artifacts but also reveals more true blood vessels, which match with those in the ground-truth images. It needs to be noted that although image-domain DNNs for motion correction are powerful in mitigating artifacts, they are less capable of revealing features not observable in the original images. In practice, for a certain region in the tissue, if most of its motions have amplitudes greater than the image resolution, the detected signals from this region will have severe mismatches, which causes feature loss in the reconstructed images. The fact that the proposed hybrid method reveals these features proves its high tolerance to tissue motion and highlights the importance of system matrix manipulation in motion correction.

Furthermore, we use the proposed method to correct heartbeat-induced motions in human breast imaging (*N*_loc_ = 99, *N*_ele_ = 4 × 256). We acquired signals from the left breast of a volunteer with breath holding (10 s) in 4 experiments. Samples of the signals in the first two experiments are shown in **Supplementary Fig. 15a** and **b**, respectively, from which we observe the signals’ temporal shifts caused by heartbeat-induced motions. We reconstruct the human breast images in the first two experiments and compare MAPs of them in **Fig. 5**. For the first experiment, the MAPs of images reconstructed without and with motion correction are shown in **Fig. 5a1** and **a2**, respectively, with their closed-up subsets (blue-dashed and red-dotted boxes) shown in **Fig. 5a2-a3** and **b2-b3**. We observe that motion correction mitigates motion-induced artifacts (e.g., the region indicated by blue-dotted arrows in **Fig. 5a2-b2**) and reveals motion-compromised features (e.g., the blood vessel indicated by red-solid arrows in **Fig. 5a3-b3**). MAPs of the images in the second experiment without and with motion correction are shown in **Fig. 5c1–c3** and **d1–d3**, respectively. Still, motion correction recovers motion-compromised vessels, such as those indicated by blue-solid and red-solid arrows in **Fig. 5c2-d2** and **c3-d3**, respectively. Importantly, the compromised features in **Fig. 5a2-a3** are very different from those in **Fig. 5c2-c3** due to motions’ different effects in the two experiments, although the breast just deforms slightly between the two experiments. After motion correction, the features have high similarity as shown in **Fig. 5b2-b3** and **Fig. 5d2-d3**, which validates that the features recovered by motion correction are true. Image qualities in the las two experiments are also improved by motion correction, as shown in **Supplementary Fig. 15c1–c3** and **d1–d3**, respectively. We summarize the images for the four experiments in **Supplementary Video 9**, which highlights the improvements brought by motion correction.

**Fig. 5.**
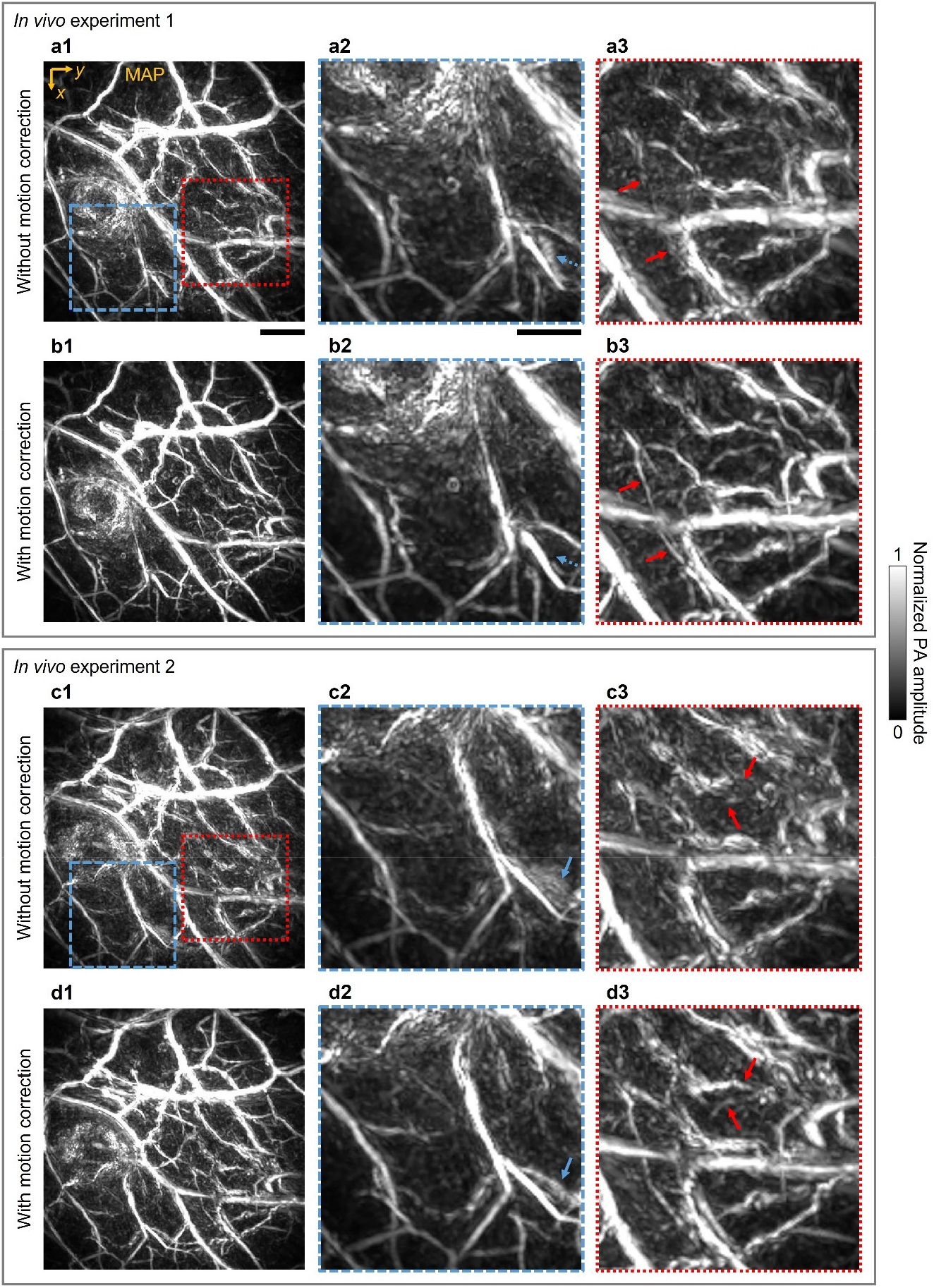
Intra-image nonrigid motion correction for human breast imaging *in vivo*. **a1–a3** MAP of a human breast image with a volume of 6 × 6 × 3 cm^3^ reconstructed without motion correction (**a1**), and two closed-up subsets marked by blue-dashed (**a2**) and red-dotted (**a3**) boxes, respectively. **b1–b3** MAP of the image reconstructed with motion correction from the same signals as for **a1**, and the closed-up subsets. Examples of suppressed artifacts and enhanced features are highlighted in **a2-b2** and **a3-b3**, respectively, by blue-dotted and red-solid arrows. **c1–c3** and **d1– d3** The second experiment of the same human breast with the images reconstructed without (**c1**– **c3**) and with (**d1–d3**) motion correction, respectively. Examples of enhanced features are indicated by blue-solid and red-solid arrows in **a2-b2** and **a3-b3**, respectively. For the first-column images, scale bar, 1 cm; for the second and third columns, scale bar, 6 mm.

## Discussion

Here, we propose a hybrid method for fast functional imaging by reconstructing images from sparsely sampled signals with a prior image and propose a method for intra-image nonrigid motion correction that does not need repeated acquisitions as required in time-gating-based methods. In both numerical simulations and mouse brain functional imaging *in vivo*, the hybrid method substantially mitigates artifacts in the reconstructed images and reduces false positive regions in the functional image, and its linearity is important for maintaining the true functional region. Due to its high robustness, the method can accelerate or enhance the performance of an existing system and reduce the cost of a future system for functional imaging. In both numerical simulations and human breast imaging *in vivo*, the motion correction method successfully mitigates motion-induced artifacts and recovers motion-compromised features. Due to its low requirement in data acquisition and high robustness, the method can be broadly applied to brain, heart, chest, abdominal, and fetus imaging. Currently, the method reduces the effects of motions in a set of signals to form a single image. In a future study, we may perform finer manipulations of the system matrix to recover the complete motion profile from these signals: an important feature for heart imaging.

Although we demonstrate the proposed methods only using PACT, they are applicable to CT and MRI. System matrices in 3D PACT and CT correspond to sphere^10^ and line^6^ integrals, respectively, and the latter can be transformed using Grangeat’s method ^45^ into plane integrals, which are locally equivalent to sphere integrals. System matrices in 2D PACT and CT^5^ correspond to circle (reduced from a sphere) and line integrals, respectively, which are locally equivalent. MRI is more complex due to its high flexibility in k-space sampling. For radial sampling MRI^35,46^, the acquired signals can be transformed to integrals on lines and planes, respectively, for 2D and 3D imaging using the Fourier slice theorem. For other sampling patterns, further analysis may disclose proper transformations to obtain integrals that are locally equivalent to those in PACT. It needs to be noted that both proposed methods rely on efficient slicing and manipulation of the system matrix, which is directly available for CT where the system matrix is highly sparse and is achieved in PACT through system matrix compression based on SVD and FFT as shown in this study. For radial sampling MRI, we can transform the k-space signals to line or plane integrals and compress them through SVD and FFT. For general MRI, further study may transform the k-space signals to compressible integrals. In summary, through certain transformations, system matrices in CT, MRI, and PACT share similar local structures, which allows the knowledge in this study to be transferred to CT and MRI.

## Methods

### System matrix compression based on SVD and FFT

We describe the original and compressed system matrices using a slow operator and a fast operator, respectively. The discretized operators are derived in **Supplementary Note 3** and summarized below:

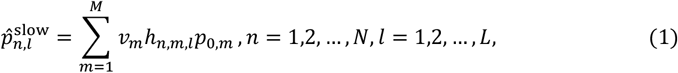

and

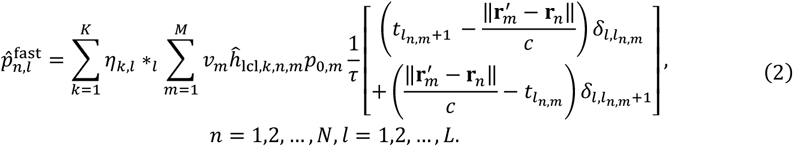

We discretize the time variable as *t*_*l*_ = *t*_1_ + (*l* − 1) *τ, l* = 1,2, …, *L*, where *t*_1_ is the initial time, *τ* is the sampling step size, and *L* is the number of time points. In the operators, 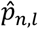 means the signal detected by the *n*-th virtual element at the time *t*_*l*_ and is approximated by 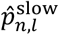 and 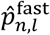, respectively; *v*_*m*_ and *p*_0,*m*_ represent the volume and initial pressure of the *m*-th voxel, respectively; *h*_*n,m,l*_ is the *n*-th virtual element’s response to the *m*-th voxel at *t*_*l*_; *η*_*k,l*_ is the *k*-th temporal singular function (obtained in SVD) evaluated at *t*_*l*_; *K* is the number of used singular value components; ∗_*l*_ means the discrete convolution for the time index *l* (implemented in FFT); *ĥ*_lcl,*k,n,m*_ is the *k*-th spatial singular function’s value depending on the *m*-th voxel’s location relative to the *n*-th virtual element; 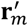 and **r**_*n*_ are the coordinates of the *m*-th voxel and *n*-th virtual element, respectively; *l*_*n,m*_ denotes the temporal index such that 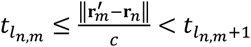, and we use the Kronecker delta function 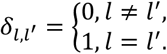. Although the numerical length of *h*_*n,m,l*_ (*l* = 1,2, …, *L*) is *L*, it has a smaller effective length, denoted as *L*′, for nonzero values. We summarize these symbols in **Supplementary Table 1**. Numerical complexities of the slow and fast operators are 0(*NML* ′) and *O*(*NMK*) + *O*(*N*(*L*log_2_*L*)*K*), respectively. Here, the big 0 notations mean that there exist constants *β*_1_, *β*_2_, and *β*_3_ (independent from *NML* ′, *NMK*, and *N*(*L*log_2_*L*)*K*, respectively) such that the computations based on the slow and fast operators, respectively, can be done in times less than *β*_1_*NML* ′ and *β*_2_*NMK* + *β*_3_*N*(*L*log_2_*L*)*K*. The slow operator consists of only spatial integration; whereas, the fast operator consists of two layers of operations, and the terms *β*_2_*NMK* and *β*_3_*N*(*L*log_2_*L*)*K* are responsible for the inner layer spatial integration with *ĥ*_lcl,*k,n,m*_ and the outer layer temporal convolution with *η*_*k,l*_, respectively. In numerical simulations with *M* = 50 × 50 × 50, *N* = 396 × 128, *L* = 4096, *L* ′ = 151, and *K* = 3 (**Supplementary Note 4**), we observe that the fast operator based on Eq. (2) has approximately 42 times the speed of the slow operator based on Eq. (1).

### A hybrid method for fast functional imaging

We first acquire signals from an object with transducers at enough locations (*N*, dense sampling). Then we continuously acquire multiple sets of signals from the object with transducers at a smaller number of locations (*N* ′, sparse sampling) for fast functional imaging. We assume that the object does not move as a whole during functional imaging.

For image reconstruction, we denote the densely sampled signals and a set of sparsely sampled signals as 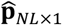 and 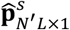, respectively. We first obtain an image 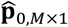 from the densely sampled signals 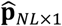 by solving the regularized optimization problem

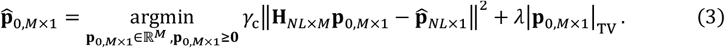

Here, *γ*_c_ represents the system-specific measurement calibration factor, meaning 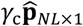 is the reading of the data acquisition system; **H**_*NL*×*M*_ is the dense-sampling system matrix; |**p**_0,*M*×1_|_TV_ denotes **p**_0,*M*×1_’s total variation (TV) norm, defined in Eq. (S35); and *λ* means the regularization parameter. To obtain an image from 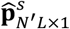 directly, we can use 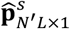 and the sparse-sampling system matrix 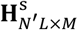 to replace 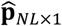 and **H**_*NL*×*M*_, respectively, in Eq. (3) and solve it. However, the nonlinearity introduced by the nonnegativity constraint and TV regularization may disrupt the functional signals. To maintain the linearity under sparse sampling, we treat 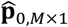 as a prior image and apply a smooth modulation to it to extract the dominant sources of aliasing artifacts by solving the optimization problem

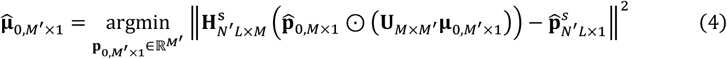

using a fast iterative shrinkage-thresholding algorithm (FISTA) with a constant step size^48^. Here, **μ**_0,*M*′ ×1_ is a modulation image in the form of a column vector of size 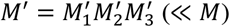, the symbol ⊙ denotes the element-wise product, and **U**_*M*×*M*′_ is an upsampling operator transferring **μ**_0,*M*′×1_ to a smooth modulation image in the column-vector form of size *M*. To implement **U**_*M*×*M*′_, we reshape the vector **μ**_0,*M*′×1_ into a 3D array 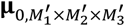, apply trilinear interpolation to the 3D array to obtain 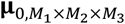, and flatten it to obtain **U**_*M*×*M*′_ **μ**_0,*M*′×1_. The array 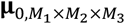 is a smooth array determined only by *M*′ independent values. Thus, the expression 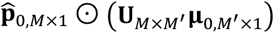 represents a smooth modulation of the prior image 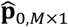. By solving the optimization problem in Eq. (4) we obtain 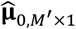 and represent the dominant sources causing aliasing artifacts by 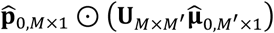, which correspond to the dominant signals causing aliasing artifacts 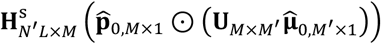. It needs to be noted that the implementation of the forward operator based on Eq. (S30) allows for efficient slicing of the system matrix, such as the matrix 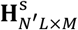, a slicing with respect to transducer element indices. Also, for a given set of parameters in FISTA, the solution to Eq. (4), 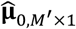, depends linearly on 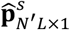.

Then, we apply the dense-sampling system matrix **H**_*NL*×*M*_ to 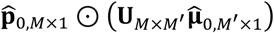 to simulate the densely sampled signals 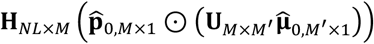, from which we use the UBP method to reconstruct an image, denoted as 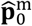. We further remove the aliasing artifacts caused by signals from 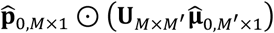 by subtracting these signals from 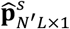 to obtain the residual signals 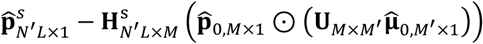 and using them with the UBP method to reconstruct an image, denoted as 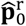. Due to the linearity of the UBP method, both images 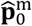 and 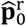 are linearly dependent on 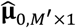 and thus on 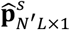. We combine these two images to obtain the final image

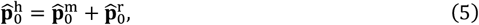

for 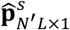. We summarize the workflow of the hybrid method in **Supplementary Fig. 6**. We repeat the process for other sets of sparsely sampled raw data to obtain the images for further functional signal extraction. Additional symbols for fast functional imaging are shown in **Supplementary Table 2**.

### Functional signal extraction

Given a set of reconstructed images 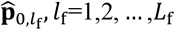 in functional imaging, we propose a method to extract functional signals from them through regularized correlation. To explain the process, we define 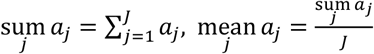, and 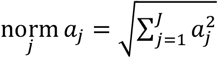 for a series of numbers *a*_*j*_, *j* = 1,2, …, *J*. We assume that the functional signal has a profile 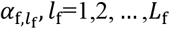, normalized to 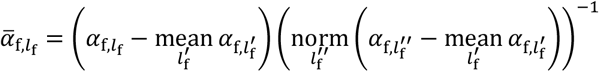. The *m*-th voxel has a value of 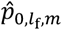 in the *l*_f_-th image 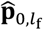. We can quantify the functional amplitude at the *m*-th voxel through the Pearson correlation coefficient (PCC) between 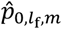 and 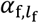:

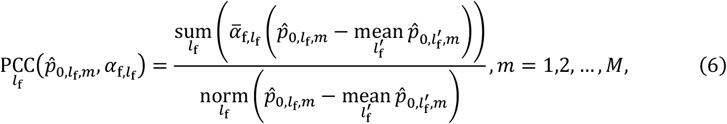

which is not robust for very-low-amplitude regions. To improve the robustness, we add a regularization term to the denominator and obtain the regularized PCC (PCCR):

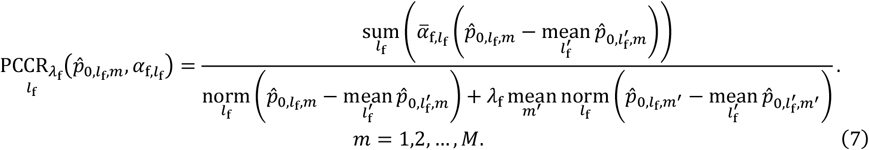

A 3D functional image is formed by assigning the regularized correlation to each voxel.

The assumed functional signal profile 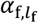 is directly available in the numerical simulations (Eq. (S37)). For mouse brain functional imaging *in vivo*, we first let 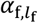 be a sinusoidal profile synchronized with the paw stimulation pattern and apply it to the UBP reconstructed images with 4*N*_loc_ = 76 to obtain the functional region. Then we extract the true functional profile from this region and apply it to images with 4*N*_loc_ = 40, 20, 12.

### Intra-image nonrigid motion correction

For intra-image nonrigid motion correction, we incorporate motion parameters into the system matrix and obtain the motions and image simultaneously by solving a dual-objective optimization problem. To describe the intra-image motion, we denote the number of elements of the four-arc array as *N*_ele_ and the number of locations of the array in a 3D imaging process as *N*_loc_. Also, we decompose the whole image domain *D* into rectangular-cuboid subdomains 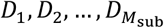 satisfying Eq. (S39). We ignore the deformations and rotations of each relatively small subdomain and only discretize the translations with step sizes *a*_*x*_, *a*_*y*_, and *a*_*z*_ along the *x*-axis, *y*-axis, and *z*-axis, respectively. We represent the translation steps of the subdomain of 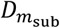 (centered at 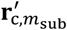) as 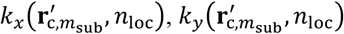 and 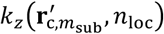 when the array moves to the *N*_loc_ -th location. Further, we define 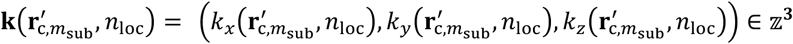 and 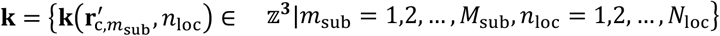, which fully characterizes the intra-image motion in 3D imaging. Incorporating **k** into the system matrix using Eqs. (S40) and (S41), we obtain **H**_**k**_. Then we express the motion correction as a dual-objective optimization problem

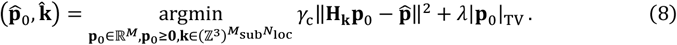

To solve this problem, we decompose it into multiple subproblems (Eqs. (S44), (S50), (S51), and (S52)) and solve them sequentially in multiple rounds of Gauss-Seidel-type iterations. A detailed discussion and the workflow of the motion correction method are shown in **Supplementary Note 8** and **Supplementary Fig. 14**, respectively, and the additional symbols are shown in **Supplementary Table 3**. The key idea is to reconstruct an initial image without motion correction, decompose the image into sub-images, estimate the motion of each sub-image for every rotating location of the four-arc array, incorporate the motion information into the system matrix for another reconstruction, and iterate the correction-reconstruction process to finalize the image.

### Numerical simulations of intra-image motions

To simulate translation- and deformation-induced motions, we let the values at 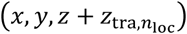 and 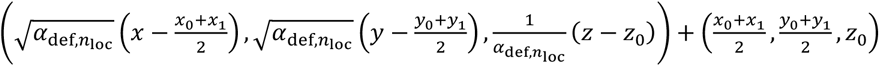 respectively, in the *n*_loc_ -th numerical phantom be the value at (*x, y, z*) ∈ [*x*_0_, *x*_1_] × [*y*_0_, *y*_1_] × [*z*_0_, *z*_1_] in a predefined numerical phantom, with the translation distances defined as

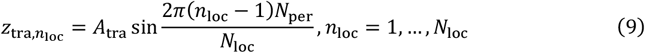

and deformation ratios defined as

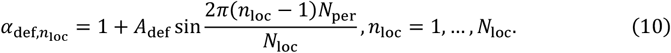

Here, *A*_tra_ and *A*_def_ are amplitudes of the translation distance and deformation ratio, respectively. In the forward simulations, signals from the *n*_loc_-th numerical phantom are only detected by the transducer array at the *n*_loc_-th location. We let *x*_1_ − *x*_0_ = 44.8 mm, *y*_1_ − *y*_0_ = 44.8 mm, *z*_1_ − *z*_0_ = 14.4 mm, *N*_loc_ = 72, *N*_ele_ = 4 × 72, and *N*_per_ = 0.5, 1, 2, 3 for all motion simulations, and use *A*_tra_ = 0.2, 0.4, 0.6, 0.8, 1.0, 1.2 (mm) and *A*_def_ = 0.02, 0.04, 0.06, 0.08, 0.10, 0.12, respectively, for translation and deformation simulations.

### Imaging protocols

The animal experiments followed the protocol approved by the Institutional Animal Care and Use Committee of the California Institute of Technology. The human imaging experiment followed the protocol approved by the Institutional Review Board of the California Institute of Technology.

## Data availability

All data used in this study are available from the corresponding author upon reasonable request.

## Code availability

The image reconstruction and data processing algorithms are described in detail in the Methods and Supplementary Information. We have opted not to make the codes available because they are proprietary and used for other projects.

## Acknowledgments

This work was supported in part by National Institutes of Health grants R01 NS102213, U01 EB029823 (BRAIN Initiative), and R35 CA220436 (Outstanding Investigator Award).

## Author contributions

P.H. and L.V.W. conceived and designed the study. X.T. and L.L. performed the experiments. P.H. implemented the image reconstruction and processing algorithms. P.H. and X.T. analyzed the data. L.V.W. supervised the study. All authors wrote the manuscript.

## Supplementary Information

### Supplementary Note 1 A fast forward operator based on singular value decomposition (SVD) and fast Fourier transform (FFT)

In a homogeneous medium, a photoacoustic wave can be expressed as^10,47^

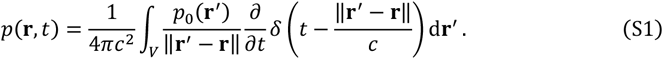

Here, *p*(**r**, *t*) is the pressure at location **r** and time *t, c* is the speed of sound (SOS), *V* is the volumetric space occupied by the tissue, and *p*_0_(**r**′) is the initial pressure at **r**′. Assume that we have *N* finite-size ultrasonic transducer elements. For the *n*-th transducer element at **r**_*n*_ with detection surface *S*_*n*_, the average pressure on the detection surface at time *t* is expressed as

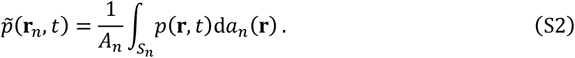

Here, *A*_*n*_ denotes the area of the surface *S*_*n*_. We denote the spatial impulse response (SIR) in Eq. (S2) as^10^

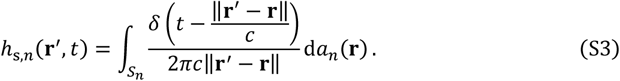

Then Eq. (S2) becomes

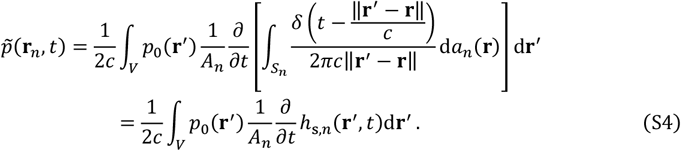

We denote the electric impulse response (EIR) of the *n*-th transducer as *h*_e,*n*_(*t*) and express the transducer’s response using temporal convolution ∗_*t*_ as

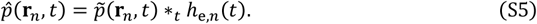

Substituting Eq. (S4) into Eq. (S5) yields

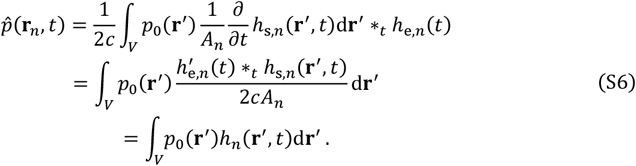

Here, the prime in 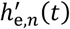 denotes the time derivative, and *h*_*n*_ denotes the point source response per unit initial pressure per unit infinitesimal tissue volume received by a finite-size transducer element:

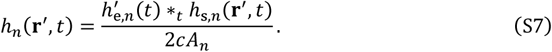

For convenience in the following discussion, we temporally shift *h*_*s,n*_(**r**′, *t*) and *h*_*n*_(**r′**, *t*) for **r**′ such that time 0 aligns with the onset of the nonzero signal received by the center of the *n*-th transducer element **r**_*n*_:

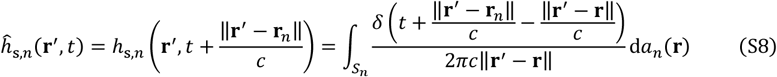

and

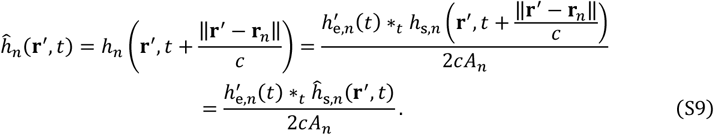

Next, we express the SIR and point source response in the local coordinates of the transducer elements. Each transducer element used in this research has a flat rectangular detection surface with a length *a* of 0.7 mm and a width b of 0.6 mm, yielding *A*_*n*_ = *ab* = 0.42 mm^2^, *n* = 1,2, …, *N*. These transducer elements also have the same EIR: *h*_e,*n*_(*t*) = *h*_e_(*t*), *n* = 1,2, …, *N*, and the measurement of *h*_e_(*t*) will be discussed in **Supplementary Note 2**. We define the local coordinates of the *n*-th transducer element using the center the detection surface as the origin, the length direction as the *x*-axis, the width direction as the *y*-axis, and the normal direction (toward the detection region) as the *z*-axis. Here, we choose the *x*-axis and *y*-axis to let the coordinates satisfy the right-hand rule. We express the three axes of the local coordinates as three vectors of unit length: 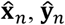, and 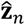 (shown in **Supplementary Fig. 1a**), which form an orthonormal matrix

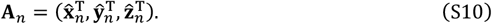

Locations **r**′, **r**_*n*_, and **r** in the global coordinates correspond to the locations 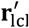, **0**, and **r**_lcl_ in the local coordinates of the *n* -th transducer element. Coordinate transformations yield 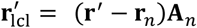 and **r**_lcl_ = (**r** − **r**_*n*_)***A***_*n*_. These global and local coordinates satisfy 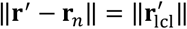 and 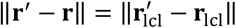 due to the orthonormality of the transformations. We denote the detection surface (*S*_*n*_ in the global coordinates) in the local coordinates as *S*_*n*,lcl_, and denote the local Leibniz notation as d*a*_*n*,lcl_(**r**_lcl_) = d*a*_*n*_(**r**) . Thus, in the local coordinates of the *n* -th transducer element, we express Eq. (S8) as

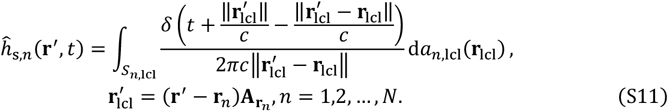

All transducer elements are geometrically identical and have the same local coordinates, meaning *S*_*n*,lcl_ = *S*_*n*′,lcl_ and d*a*_*n*,lcl_ (**r**_lcl_) = d*a*_*n*′,lcl_(**r**_lcl_) for *n, n*′ ∈ {1,2, …, *N*} . We define *S*_lcl_ = *S*_1,lcl_, d*a*_lcl_ (**r**_lcl_) = d*a*_1,lcl_(**r**_lcl_), and rewrite Eq. (S11) as

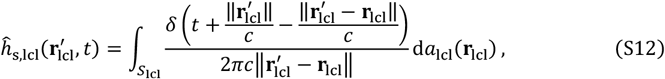

which is now independent of the transducer element index *n*. Replacing 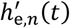 and ĥ_*s,n*_(**r**^**r**^, *t*) with 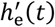 and 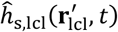, respectively, in Eq. (S9), we define

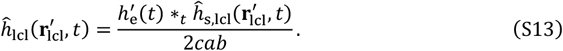

Thus, we need to calculate only the values of 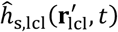 and 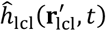, then obtain the values of *ĥ*_*s,n*_(**r**′, *t*) and *ĥ*_*n*_(**r**′, *t*) through coordinate transformation:

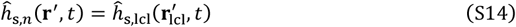

and

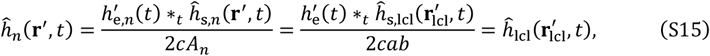

respectively, with 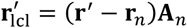 for *n* = 1,2, …, *N*. Through these relations, we express both the SIR and the point source response in the local coordinates.

**Supplementary Fig. 1.**
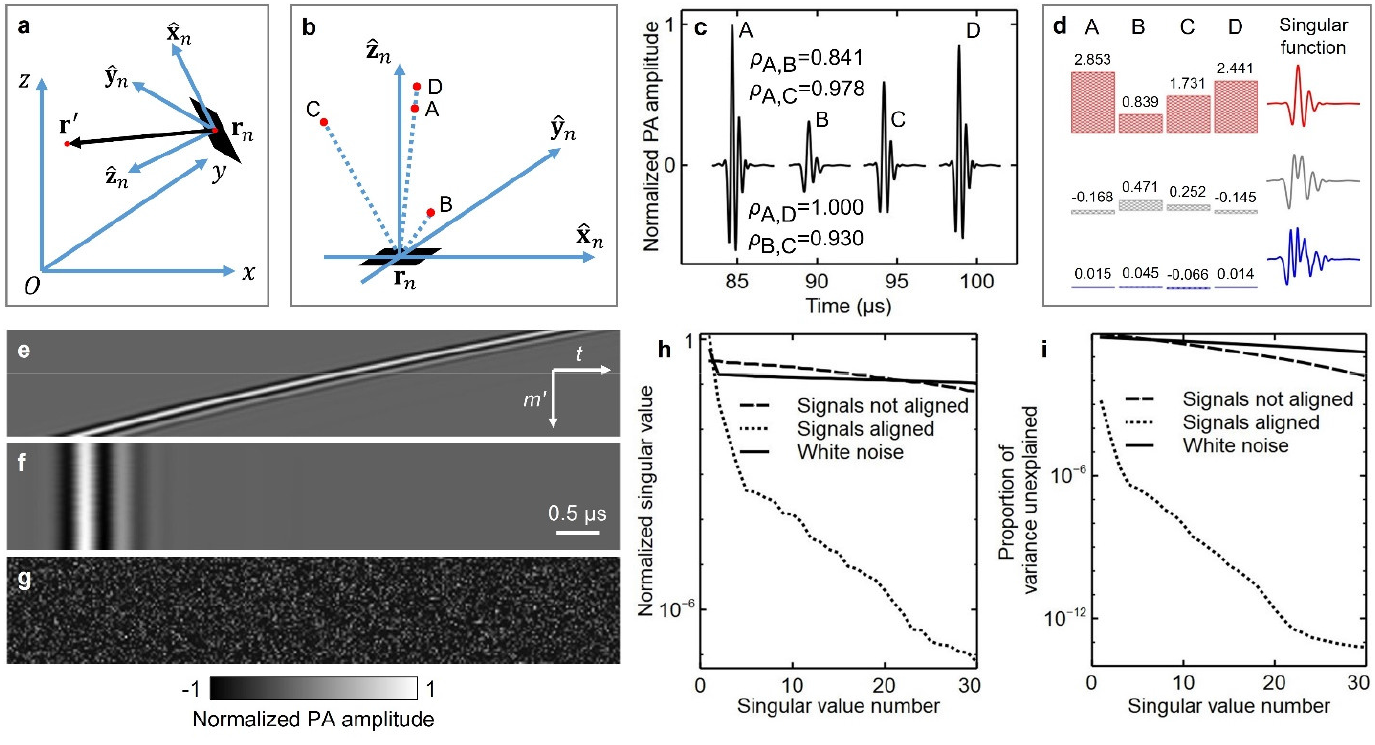
Compression of the forward operator based on SVD. **a** A point source (a red dot), the *n*-th transducer element (a black rectangular centered at **r**_*n*_), and the element’s local coordinate system with axes 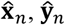, and 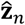. **b** Four point sources A, B, C, and D (red dots) in the local coordinate system of the *n*-th transducer element. Points A, D, and **r**_*n*_ are on the same line. **c** The responses of the transducer element to the signals from the four point sources and the PCC between every two responses. **d** Expression of the four responses using linear combinations (coefficients visualized with bars) of three temporal singular functions shown as red, gray, and blue curves, respectively, based on SVD. **e-f** Independent responses (**e**) of a transducer element to 50 point sources with decreasing distances to the element, and the temporally-shifted form (**f**), which aligns the nonzero signals in time. **g** White-noise responses of the same size as those in **e** and **f. h-i** Normalized singular values and proportions of the variances unexplained, respectively, in the SVDs of the signals in **e, f**, and **g**.

We visualize the signals detected by a transducer element by picking four point sources, labeled as A, B, C, and D, respectively, in the local coordinate system of the *n*-th transducer element (**Supplementary Fig. 1b**), with points A, D, and the element center **r**_*n*_ on the same line. The element’s responses to the signals from the point sources are shown in **Supplementary Fig. 1c**. We let *ρ*_*A,B*_ denote the Pearson correlation coefficient (PCC) between the responses corresponding to *A* and B. A direct implementation of the forward operator based on Eq. (S6) is computationally intensive. Although *ρ*_*A,D*_ = 1, due to the effects of SIR, *ρ*_*A,B*_, *ρ*_*A,C*_, and *ρ*_*B,C*_ are less than 1, indicating the signals from points A, B, and C are not shift-invariant; thus, an efficient temporal convolution with one kernel function cannot yield the detected signals accurately.

To accelerate the forward operator, we decouple the spatial and temporal dimensions of 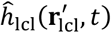 using singular value decomposition (SVD) while keeping only the dominant components:

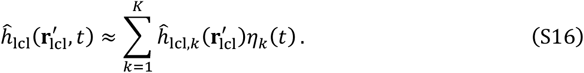

Here, 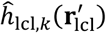 and *η*_*k*_ denote the *k*-th spatial singular function and the *k*-th temporal singular function, respectively; and we use the first *K* terms to approximate the whole series. Combining Eqs. (S9), (S15), and (S16), we obtain

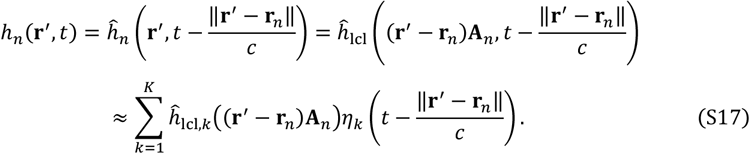

Substituting Eq. (S17) into Eq. (S6), we obtain

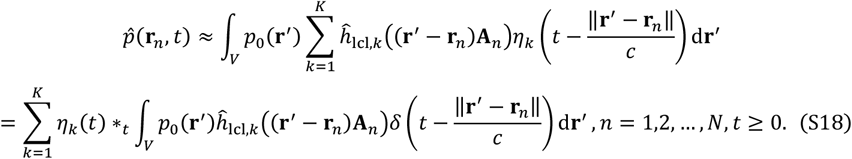

As shown in this equation, we split the temporal variable from the spatial integrals, which allows for a fast implementation of the forward operator.

We apply Eq. (S16) with *K* = 3 to the responses in **Supplementary Fig. 1c** for an initial demonstration. The three temporal singular functions *η*_1_(*t*), *η*_2_(*t*), *η*_3_(*t*) are shown as red, gray, and blue curves, respectively, in **Supplementary Fig. 1d**, and the values of spatial singular functions are visualized as bars. Then we explain the necessity of temporal shifting for alignment, described in Eqs. (S8) and (S9), for SVD. We select 50 point sources with decreasing distances to a transducer element and visualize the element’s independent responses to them in **Supplementary Fig. 1e**. The temporally-shifted form of these responses based on Eq. (S9) is shown in **Supplementary Fig. 1f**. We also add white-noise responses with the same size for comparison, as shown in **Supplementary Fig. 1g**. Performing SVD to the three sets of responses, we observe different compression efficiencies from the perspectives of normalized singular value (**Supplementary Fig. 1h**) and proportion of the variance (**Supplementary Fig. 1i**). In both figures, we see that the compression efficiency of the original responses is similar to that of the white-noise responses, whereas the efficiency for the temporally-shifted responses is significantly higher (necessary for the compression).

### Supplementary Note 2 Point source response of an ultrasonic transducer with a flat rectangular detection surface

The fast operator can be configured for a transducer with any detection surface. Here, we only discuss an ultrasonic transducer with a flat rectangular detection surface, which is used in our 3D imaging system^43^. We see from Eqs. (S6) and (S15) that the forward operator is based on the point source response 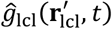 in the local coordinates. The response is determined by the first derivative of 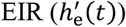 and the realigned 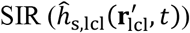 in the local coordinates. In general, we can measure 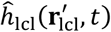 experimentally. In this research, for an ultrasonic transducer with a flat rectangular detection surface, we calculate 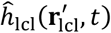 efficiently using the far-field approximation of 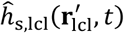^10^. From a special case of this approximation, we derive a method to measure 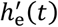 experimentally.

In the first step of obtaining 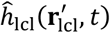, we introduce the calculation of 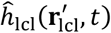 based on the far-field approximation of 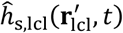. In this research, the image domain is within the far-field regions of all transducer elements. Using the far-field approximation, we express the SIR in the frequency domain as^10^

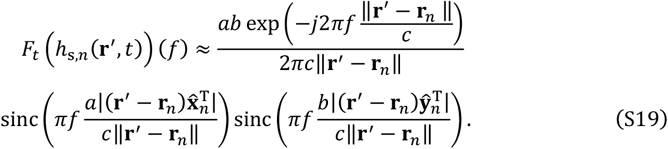

Here, *F*_*t*_ denotes the temporal FT, and *j* denotes the imaginary unit. Combining Eqs. (S14) and (S19), we apply the temporal Fourier transform (FT) to 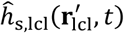 and obtain

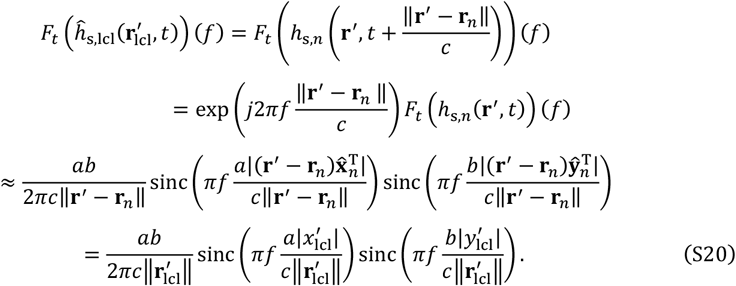

Here, we use identities 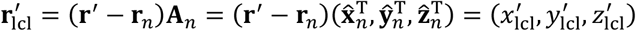 and 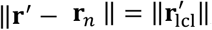, and 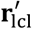 must not be the origin in the local coordinates. Substituting Eq. (S20) into the temporal FT of Eq. (S13), we obtain

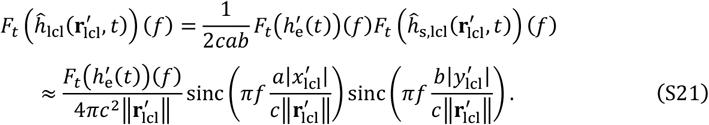

Applying temporal inverse FT to Eq. (S21), we obtain

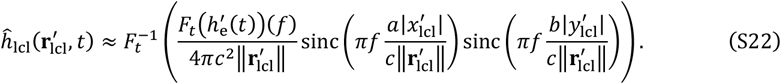

In the second step of obtaining 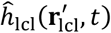, we derive a method to measure 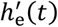 by analyzing a special case of Eq. (S22). We constrain the location 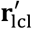 on the axis of the transducer by letting 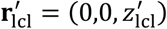, which simplifies Eq. (S22) to

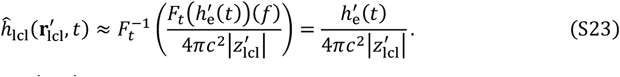

Solving for 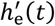 from Eq. (S23), we obtain

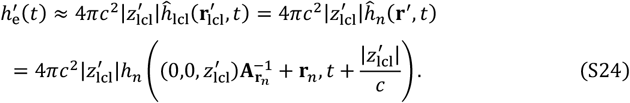

Here, we use the identity 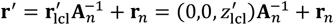. In practice, we repeat the measurement of the right-hand side of Eq. (S24) and use the average to represent 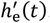.

In summary, we measure 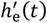 experimentally on the basis of Eq. (S24) and substitute the measurement into Eq. (S22) to obtain 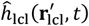. Further, we perform SVD to 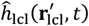 according to Eq. (S16) and obtain singular functions 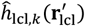 and *η*_*k*_(*t*), *k* = 1,2, …, *K*. We use these functions in Eq. (S18) to implement the fast forward operator.

### Supplementary Note 3 Discretization of the forward operator

We express the forward operator in two different forms in Eqs. (S6) and (S18), respectively. First, we discretize the forward operator implemented in Eq. (S6). In the temporal domain, we choose points of interest *t*_*l*_ = *t*_1_ + (*l* − 1) *τ, l* = 1,2, …, *L*, where *t*_1_ is the initial time, τ is the sampling step size, and *L* is the number of time points. Then we discretize the temporal FT of 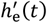 as 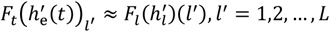, where we define 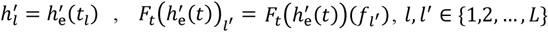, and let *F*_*l*_ represent the discrete FT with respect to *l* . The temporal frequencies f_*l*′_, *l*′ = 1,2, …, *L* are selected according to the requirement of the discrete FT. We further denote the *m*_lcl_ -th location in the local coordinates as 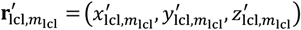 and define 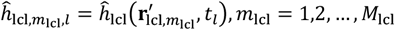, where *M*_lcl_ is the number of locations of interest in the local coordinates. Thus, we discretize Eq. (S22) as

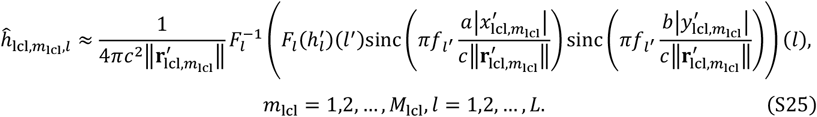

Further, we discretize the point source response *h*_*n*_(**r**′, *t*) as 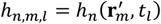, *n* = 1,2, …, *N, m* = 1,2, …, *M, l* = 1,2, …, *L*. Here, *M* is the number of voxels (source points) in the image domain. On the basis of the relation 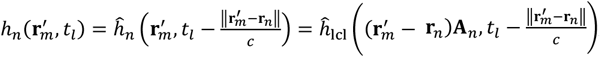, we obtain the values of *h*_*n,m,l*_ through spatiotemporal interpolation of the values of 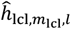. Denoting 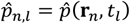 and 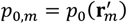, we discretize Eq. (S6) as

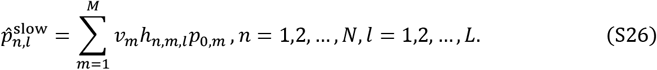

Here, 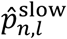 represents an approximation of 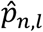 using this relatively slow operator, and *v*_*m*_ represents the volume of the *m*-th voxel. In practice, due to the finite duration of every point source response (shown in **Supplementary Fig. 1c, e**, and **f**), given a combination of *m* and *n*, we need to calculate only for *l* in a range of length *L*′ < *L*. Here, *L*′ is the effective length for nonzero values in the discretized point source responses, and the computational complexity of the discrete forward operator in Eq. (S26) is *O*(*NML*′).

Next, we discretize the forward operator to a form with lower computational complexity from Eq. (S18) . We denote 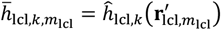 and *η*_*k,l*_ = *η*_*k*_(*t*_*l*_), *k* = 1,2, …, *K, m*_lcl_ = 1,2, …, *M*_lcl_, *l* = 1,2, …, *L*. After obtaining the array 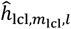 through Eq. (S25), we estimate 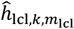 and *η*_*k,l*_ through SVD:

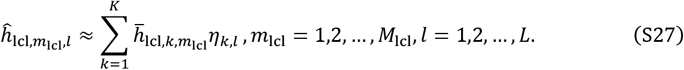

We denote 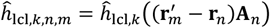, which is obtained from the values of 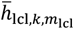 through spatial interpolation. Moreover, we express 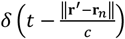 in the discrete form as

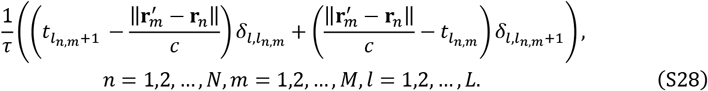

Here, *l*_*n,m*_ denotes the temporal index such that 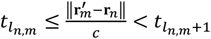, and we use the Kronecker delta function

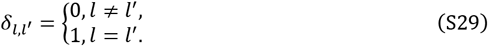

We discretize the forward operator in Eq. (S18) as

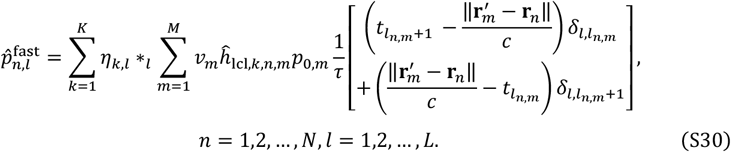

Here, 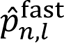 represents an approximation of 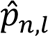 using the fast operator, and ∗_*l*_ denotes the discrete convolution with respect to *l*. In practice, we choose a value of *L* so that log_2_*L* is an integer, and we implement the discrete convolution using the temporal fast Fourier transform (FFT). Thus, the computational complexity of the discrete forward operator in Eq. (S30) is 0(*NMK*) + *O*(*N*(*L*log_2_*L*)*K*).

### Supplementary Note 4 Efficiency and signal-domain accuracy of the fast forward operator

To quantify the efficiency and accuracy of fast forward operator based on Eq. (S30), we perform numerical simulations by placing a numerical phantom of size 1 × 1 × 1 cm^3^ (**Supplementary Fig. 3a**, with voxel values of 1 inside the solid and 0 elsewhere) in four image subdomains of the system (*D*_1_, *D*_2_, *D*_3_, and *D*_4_ shown in **Supplementary Fig. 3b**). The virtual 2D array formed by the rotation of the four arc arrays is marked by blue arcs and the rotation of the four subdomains around the same axis covers a domain (marked by black-dotted circles and arcs) enclosing all the image domains discussed in this study. We also implement the slow operator according to Eq. (S26) for comparison.

We first quantify the efficiency of the fast operator by performing forward simulations with the numerical phantom in the subdomain *D*_1_ using both operators with parameters *M* = 50 × 50 × 50 (voxel size of 0.2 × 0.2 × 0.2 mm^3^), *N* = 396 × 128 (the four-arc array rotated for 99 locations and each arc downsized to 128 elements), *L* = 4096, *L*′ = 151, and *K* = 3. We ran the single-thread-CPU version of each operator 36 times on a server with Ubuntu 20.04.6 LTS and Intel(R) Xeon(R) Gold 6248R CPU @ 3.00GHz. Denoting the computation times of both operators as *t*_fast_ and *t*_slow_, respectively, we obtain me*an*(*t*_fast_) = 6.3 mi*n, st*d(*t*_fast_) = 0.4 mi*n*, me*an*(*t*_slow_) = 267 mi*n*, and *st*d(*t*_slow_) = 19 mi*n* for the 36 simulations and compare the values in **Supplementary Fig. 3c**. Further, we perform a Welch’s *t*-test between 42*t*_fast_ and *t*_slow_, showing insignificant difference (*p*-value = 0.59). Thus, the fast operator with *K* = 3 has approximately 42 times the speed of the slow operator, and the acceleration ratio can be further improved by reducing *K*.

**Supplementary Fig. 2.**
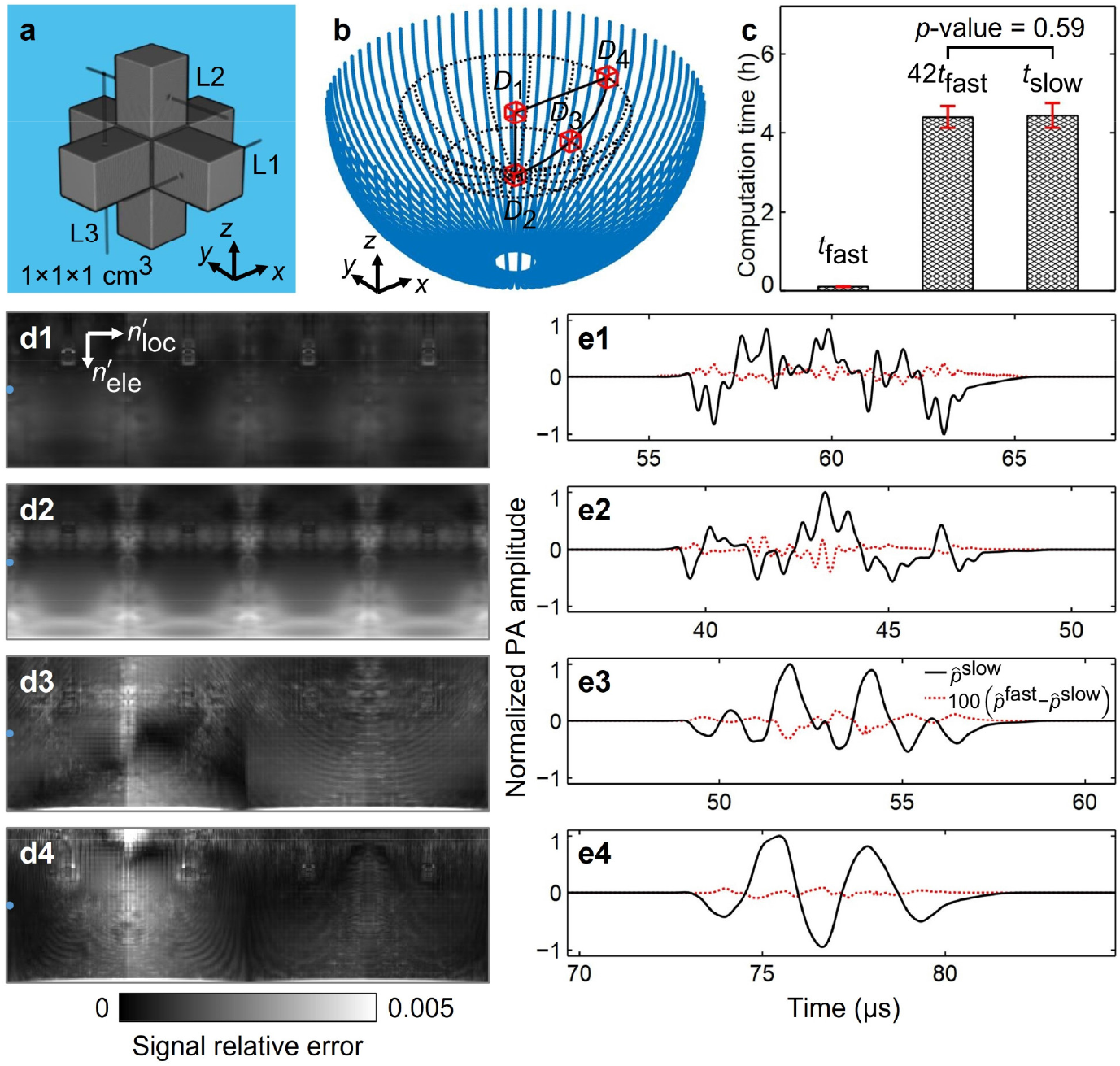
Accuracy of the fast operator in forward simulation. **a** A numerical phantom formed by three rectangular cuboids (each of size 2.6 × 2.6 × 10 mm^3^) intersecting at their centers. Voxel values are 1 inside the phantom and 0 outside. Three lines (L1, L2, and L3) are picked for image-domain comparisons **Supplementary Fig. 3. b** A virtual 2D array (blue-solid arcs), four subdomains of size 1 × 1 × 1 cm^3^ (*D*_1_, *D*_2_, *D*_3_, and *D*_4_), and the domain occupied by rotations of the four subdomains around the array axis (black-dotted circles and arcs). **c** Computation times (*t*_fast_ and *t*_slow_) of the fast and slow operators, respectively, for the forward simulations with the numerical phantom in *D*_1_ (36 repetitions). **d1–d4** Relative errors of the simulated signals for all virtual elements 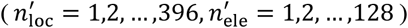, with the numerical phantom in *D*_1_, *D*_2_, *D*_3_, and *D*_4_, respectively. **e1–e4** Compares of the signals (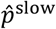 and 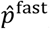) simulated by the fast and slow operators for the 64-th element 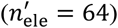 in the first virtual arc array 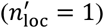, with the phantom in *D*_1_, *D*_2_, *D*_3_, and *D*_4_, respectively.

Then, we demonstrate the accuracy of the fast operator by performing forward simulations for the numerical phantom at the four subdomains (all the other parameters are the same as above). We regard the results from the slow operator as ground truth 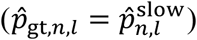 and define the relative error of the fast-operator results 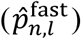 as 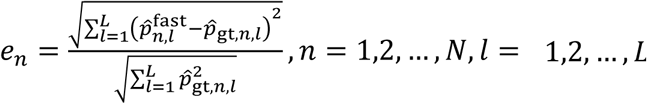. The values of e_*n*_ for the four subdomains (*D*_1_, *D*_2_, *D*_3_, and *D*_4_) are compared in **Supplementary Fig. 3d1–d4**, respectively, with 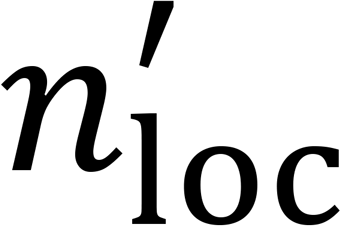 the index of the virtual arc array and *n*′ the index of the element in each array, 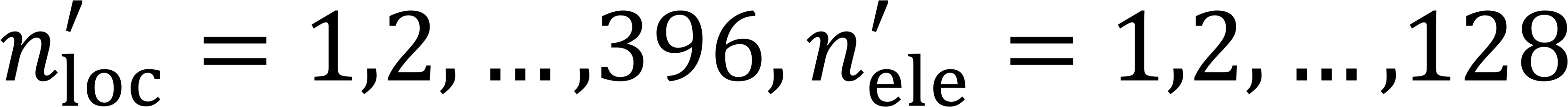. These relative errors are generally small and all of them are below 0.005, which is enough for this study. For the 64-th element 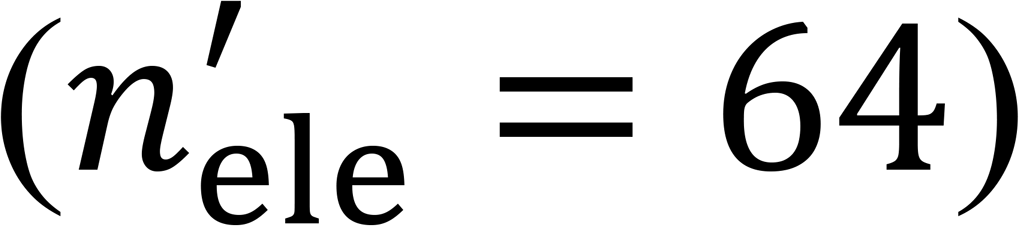 on the first virtual array 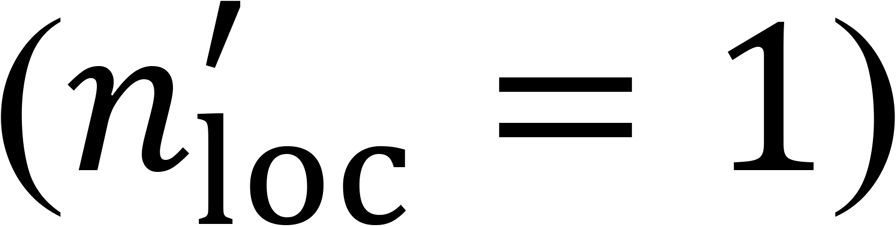, marked as blue dots in **Supplementary Fig. 3d1–d4**, we compare the signals generated by the slow and fast operators, denoted as 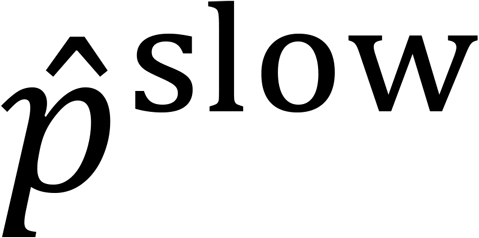 and 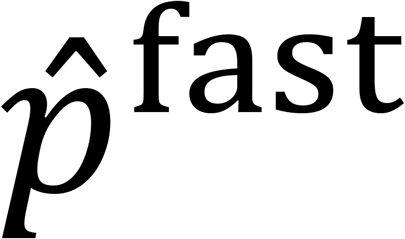, respectively, in **Supplementary Fig. 3e1–e4**, which show the fast operator’s high accuracy across the time domain.

### Supplementary Note 5 Iterative reconstruction and image-domain accuracy of the fast forward operator

We denote the discretized initial pressure and transducer response as column vectors *p*_0_ and *p*, respectively, and express the forward model as

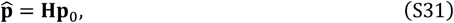

where **H** is the discretized system matrix of shape (*NL, M*). If **H** is full column rank (columns of the matrix are linearly independent), we can reconstruct an image by solving the optimization problem

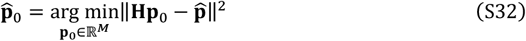

using FISTA with a constant step size^48^. We obtain the Lipschitz constant (the largest eigenvalue) of 2**H**^T^**H** through the power method with 32 iterations and use it in FISTA. Further, we can add nonnegativity of the initial pressure as a constraint to facilitate the reconstruction:

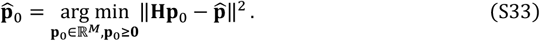

If **H** is not full column rank or the noise level is high, we reconstruct the image by solving the regularized optimization problem

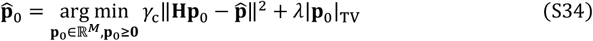

using FISTA with a constant step size^48^. Here, *λ* denotes the regularization parameter and *γ*_c_ represents the system-specific measurement calibration factor. In practice, instead of dealing with 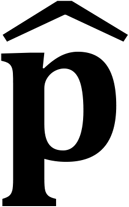 directly, we obtain 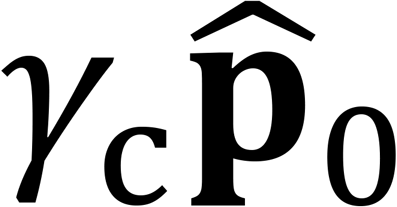 from 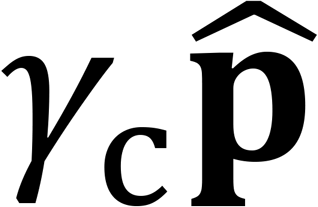 (reading of the data acquisition system) by solving the problem 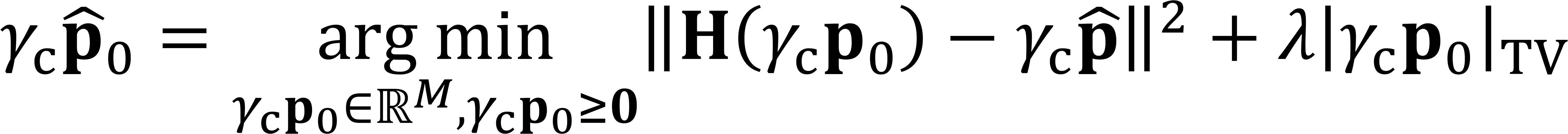, which is equivalent to Eq. (S34). The normalized values of 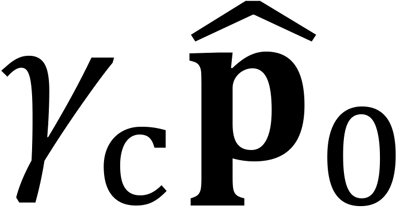 (the same as those of 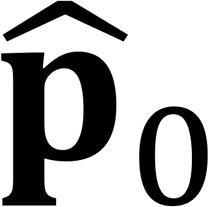) are used for further analysis, and we ignore the effect of *γ*_c_ in this study. The term |*p*_0_|_TV_ in Eq. (S34) denotes the total variation (TV) norm, defined as

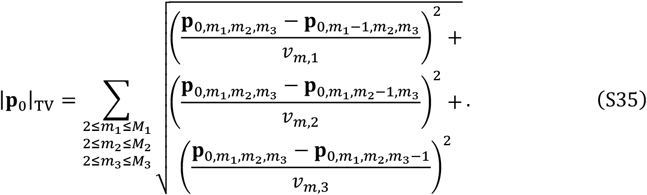

In this definition, *v*_*m*,1_, *v*_*m*,2_, and *v*_*m*,3_ represent the voxel sizes along the *x*-axis, *y*-axis, and *z*-axis, respectively; *M*_1_, *M*_2_, and *M*_3_ (*M* = *M*_1_*M*_2_*M*_3_) denote the number of voxels in the first, second, and third dimensions, respectively; and we reshape the column vector *p*_0_ to a 3D array 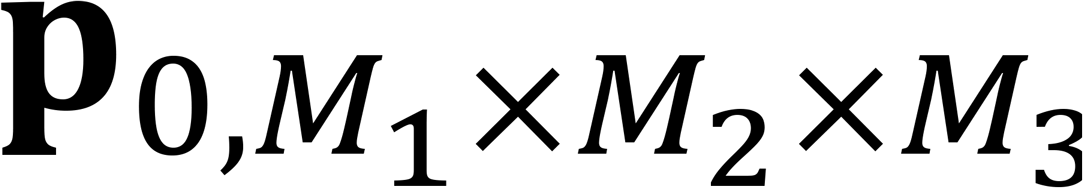 to express 3D information. Given a set of parameters for FISTA, the solution to Eq. (S32) is linearly dependent on 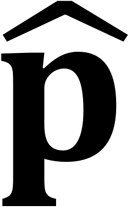. The solutions to Eqs. (S33) and (S34) are nonlinearly dependent on 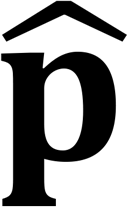 due to the nonnegative constraint (***p***_0_ ≥ **0**) and the TV regularization. Symbols for the forward operator and image reconstruction are summarized in **Supplementary Table 1**.

We quantify the accuracy of the fast forward operator in the image domain by performing forward simulations using the slow operator (**H** implemented through Eq. (S26), regarded as ground truth) and image reconstructions using iterations with nonnegativity constraints (Eq. (S33) with **H** implemented through Eq. (S30) and accelerated by two NVIDIA A100 GPUs), with the numerical phantom (**Supplementary Fig. 2a**) in the four subdomains *D*_1_, *D*_2_, *D*_3_, and *D*_4_, respectively. We define the reconstructed image 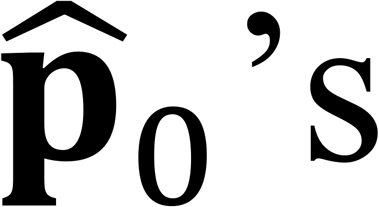 relative error to the ground-truth image ***p***_0,gt_ as 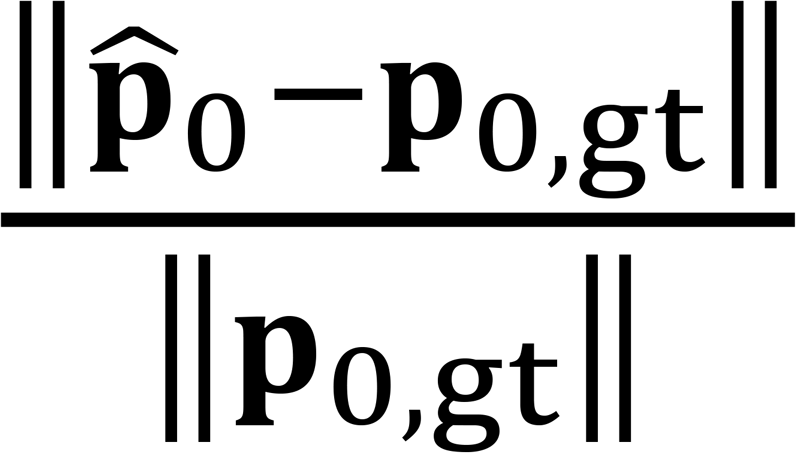. For the reconstructions with the numerical phantom in *D*_1_, the relative error reduces to 0.003 as the iteration number *n*_iter_ reaches 256, as shown in **Supplementary Fig. 3a1**. We plot the values in the reconstructed images along three lines (L1, L2, and L3 shown in **Supplementary Fig. 2a**), in **Supplementary Fig. 3a2–a4**, respectively, for *n*_iter_ = 1, 4, 8, 64, 256. Further, we denote the ground-truth (256-iteration reconstructed) values along the lines as 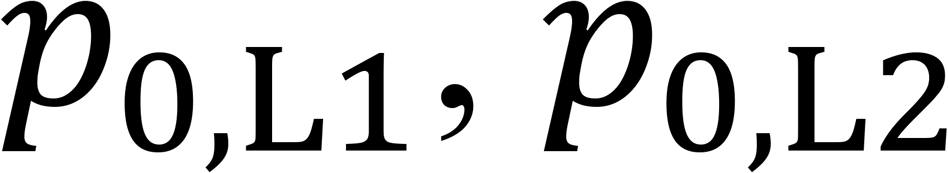, and *p*_0,*L*3_ (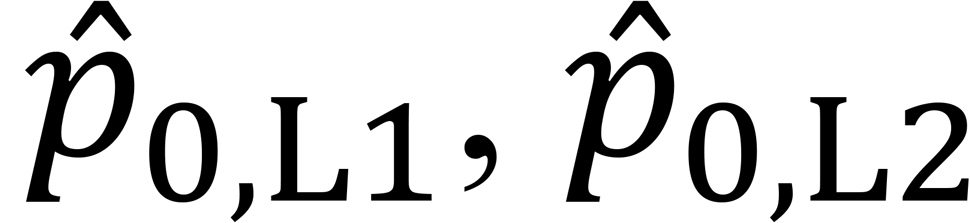, and 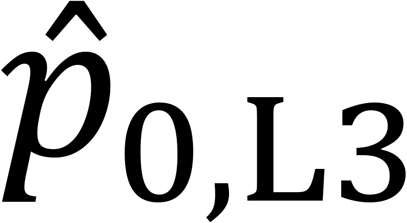), respectively, and compare them in **Supplementary Fig. 3a5**. All the plots show the high accuracy of the fast forward operator in the image domain. Repeating the analysis in subdomains *D*_2_, *D*_3_, and *D*_4_ further confirms the accuracy, as shown in **Supplementary Fig. 3b1–b5, c1–c5**, and **d1–d5**, respectively.

**Supplementary Fig. 3.**
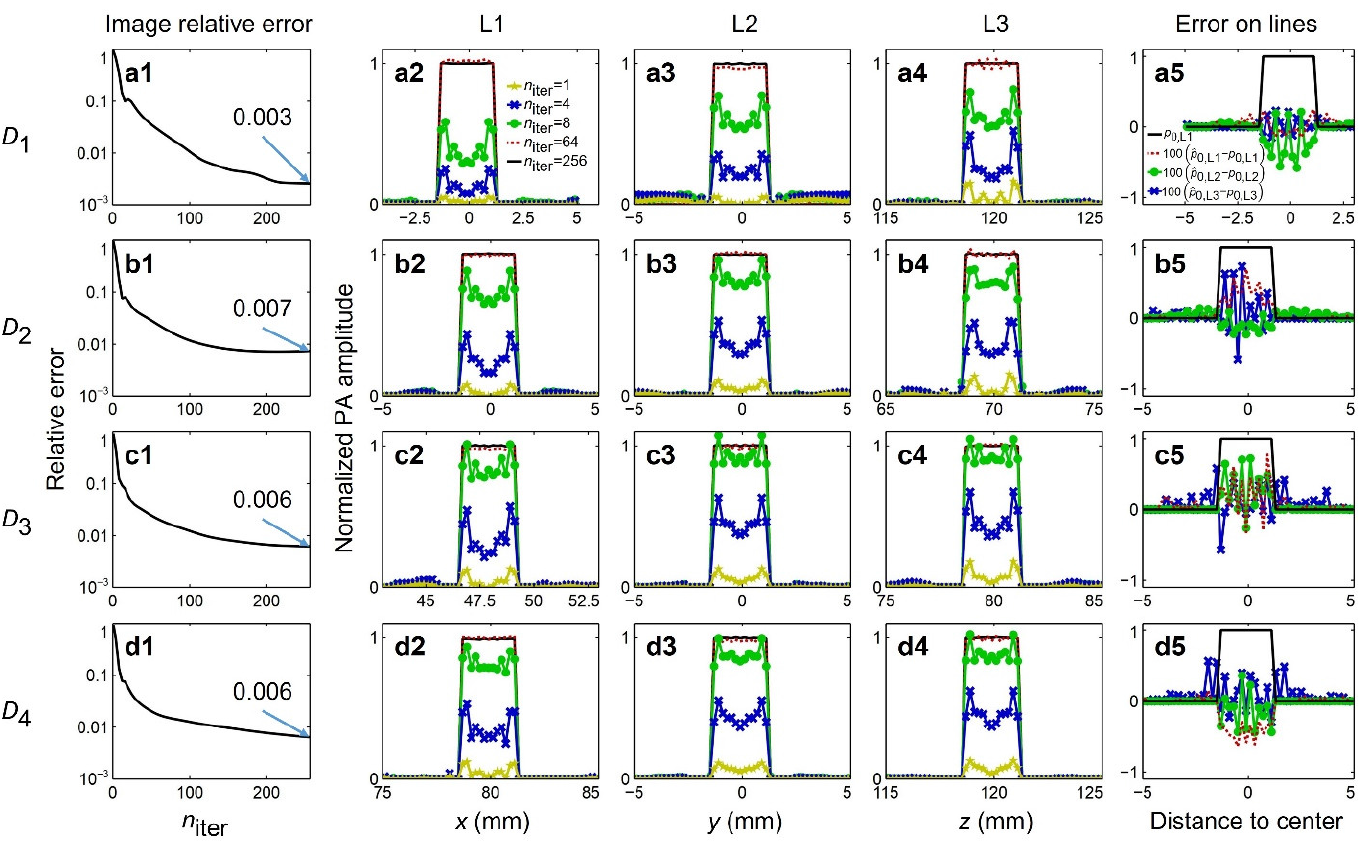
Image-domain accuracy of the fast forward operator. **a1** Relative error of the reconstructed image in the subdomain *D*_1_ with 1 to 256 iterations. **a2–a4** Values on the lines L1, L2, and L3 (shown in **Supplementary Fig. 2a**), respectively, in the reconstructed images with *n*_i*t*e**r**_ = 1, 4, 8, 64, 256. **a5** Comparison between the ground truth (*p*_0,*L*1_, *p*_0,*L*2_, and *p*_0,*L*3_) and 256-iteration reconstructed (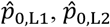, and 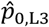) values along the three lines. **b1–b5, c1–c5**, and **d1**– **d5** The same analysis for reconstructions in subdomains *D*_2_, *D*_3_, and *D*_4_, respectively.

### Supplementary Note 6 Iterative reconstruction with noisy data or sparse sampling

After validating the accuracy of the fast forward operator in both the signal domain and image domain, we use its GPU-accelerated implementation for both forward simulation and image reconstruction in the following analyses.

First, we perform a forward simulation with the numerical phantom in the subdomain *D*_1_ (**Supplementary Fig. 2a-b**), add white noise with different amplitudes to the signals, and reconstruct the images through iterations with nonnegativity constraint (Eq. (S33)) and iterations with TV regularization (Eq. (S34)), respectively. We define the amplitude of the signal as half of the difference between its maximum and minimum values, the amplitude of noise as its standard deviation, and their division as signal-to-noise ratio (SNR). We still use the relative error 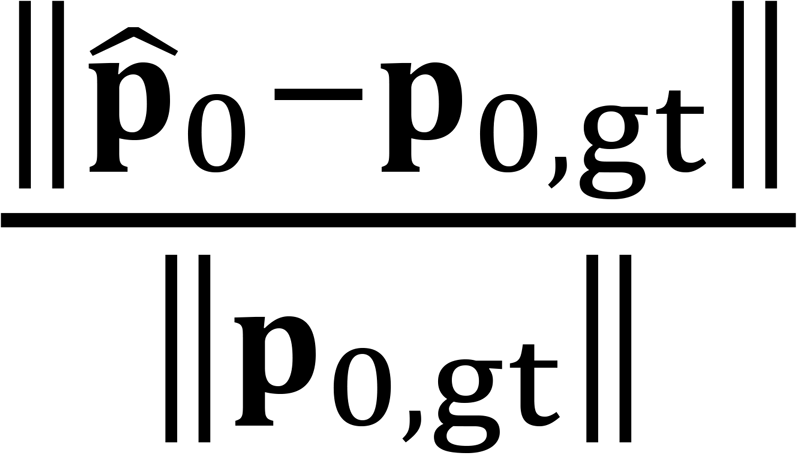 between the reconstructed image 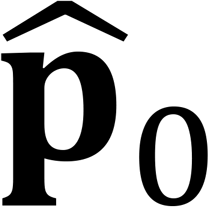 and ground truth *p*_0,gt_ to quantify the accuracy. For SNRs of 2.4, 1.2, 0.8, and 0.6, the relative errors of the images reconstructed without TV regularization are shown in **Supplementary Fig. 4a**, which means that the reconstructions are unsuccessful. For iterations with TV regularization, the relative errors for regularization parameters of 5.0 × 10^5^, 1.0 × 10^6^, 2.0 × 10^6^, and 4.0 × 10^6^ (mm) are shown in **Supplementary Fig. 4b1–b4** for the four SNRs, respectively, which demonstrate that TV regularization stabilizes the iterations and the best choice of regularization parameter (enclosed in black-solid boxes) is noise level dependent. Plots with the best regularization parameters in **Supplementary Fig. 4b1–b4** are summarized in **Supplementary Fig. 4c**, which satisfies that the higher the signal noise level the poorer the image quality. Values on the lines L1, L2, and L3 (**Supplementary Fig. 2a**) in the reconstructed images after 8, 32, and 256 iterations are shown in **Supplementary Fig. 4d1–d3, e1–e3**, and **f1–f3**, respectively. The fact that 8 iterations already reconstruct a large portion of the image inspires us to perform only 8 iterations in certain steps of the motion correction discussed below, where the general structure is more important than accurate amplitude. The fact that 256 iterations bring minor improvements to 32 iterations inspires us to perform only 32 iterations for image reconstruction in functional imaging and the final step of motion correction as discussed below.

**Supplementary Fig. 4.**
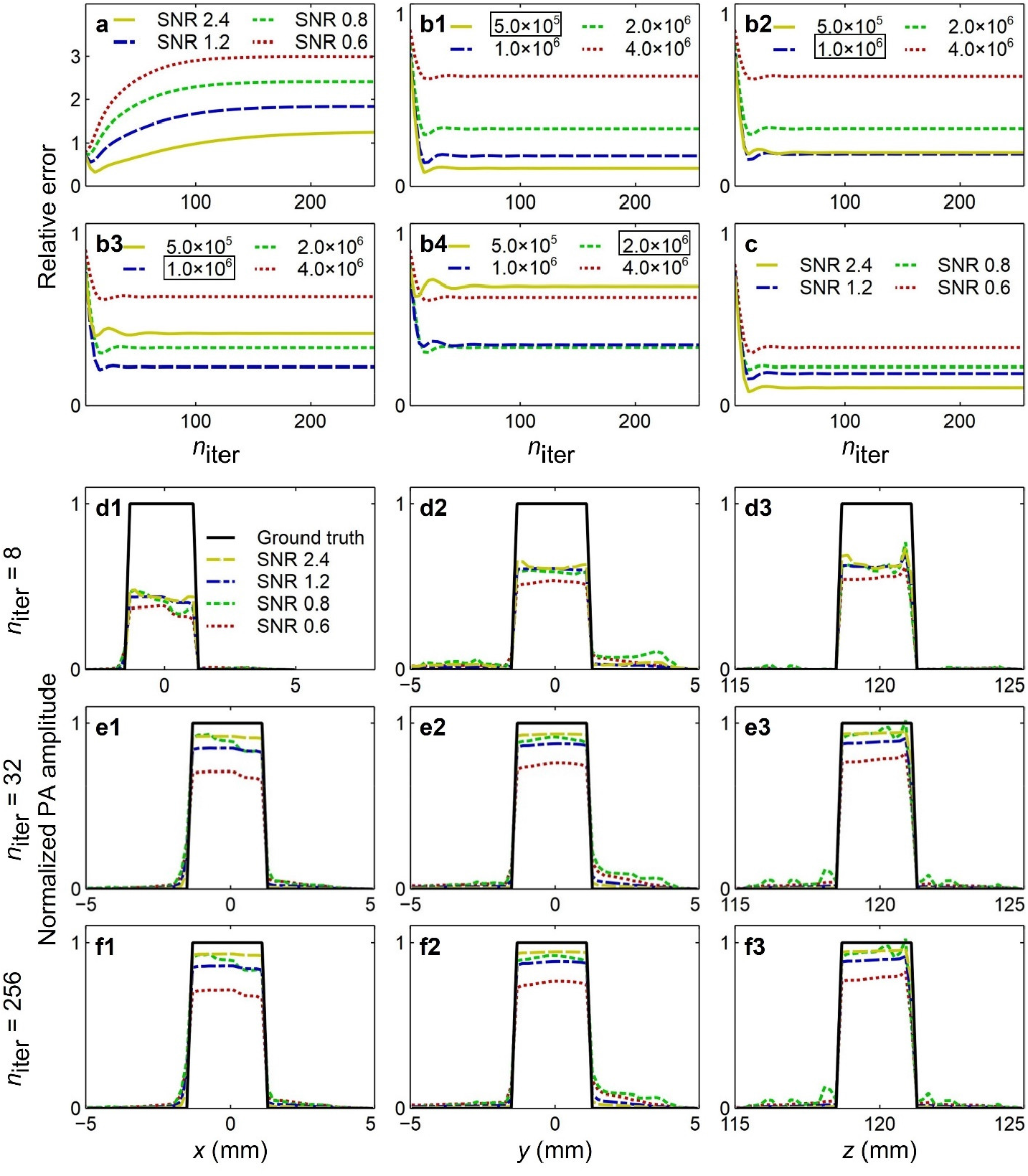
Iterative reconstruction for noisy data. **a** Relative errors of the images of the numerical phantom in *D*_1_ reconstructed iteratively with nonnegativity constraint, for SNRs of 2.4, 1.2, 0.8, and 0.6. **b1–b4** Relative errors of the images reconstructed iteratively with TV regularization, for SNRs of 2.4, 1.2, 0.8, and 0.6, respectively, for different choices of regularization parameters (mm). **c** A summary of the plots in **b1–b4** with the best choices of regularization parameters (enclosed in black-solid boxes). **d1–d3, e1–e3**, and **f1–f3** Values on the lines L1, L2, and L3 in the images reconstructed with 8, 32, and 256, iterations, respectively.

Then we use a regularized iterative method to reconstruct a mouse brain numerical phantom with signals from the phantom detected by virtual arrays with different numbers of arcs: 4*N*_loc_ = 76,40,28,20,16,12, as shown in the fourth column in **Fig. 2a**. Maximum amplitude projections (MAPs) of the images (along the *z*-axis) reconstructed by regularized iterative method with different regularization parameters for 4*N*_loc_ = 76, 40, 28, 20, 16, 12 are shown in **Supplementary Fig. 5**. Here, instead of selecting the regularization parameters quantitatively according to the relative error as in **Supplementary Fig. 4b1–b4**, we select them qualitatively by balancing mitigating artifacts and maintaining image resolution, which mimics the reality where the ground truth is unknown. The best images are indicated by red-solid boxes in **Supplementary Fig. 5**.

**Supplementary Fig. 5.**
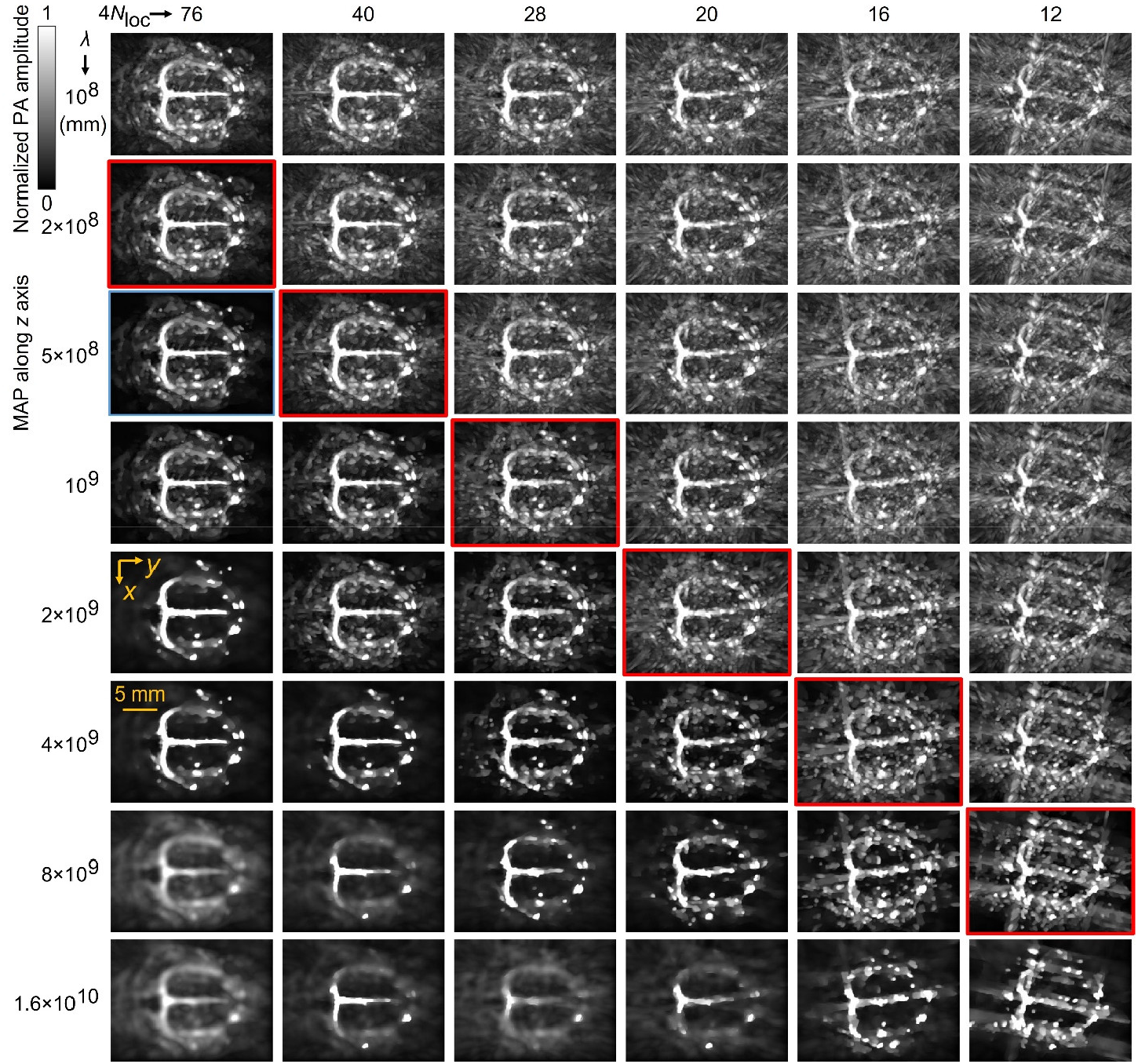
Regularized iterative reconstruction of mouse brain images with sparse sampling.

**Supplementary Fig. 6.**
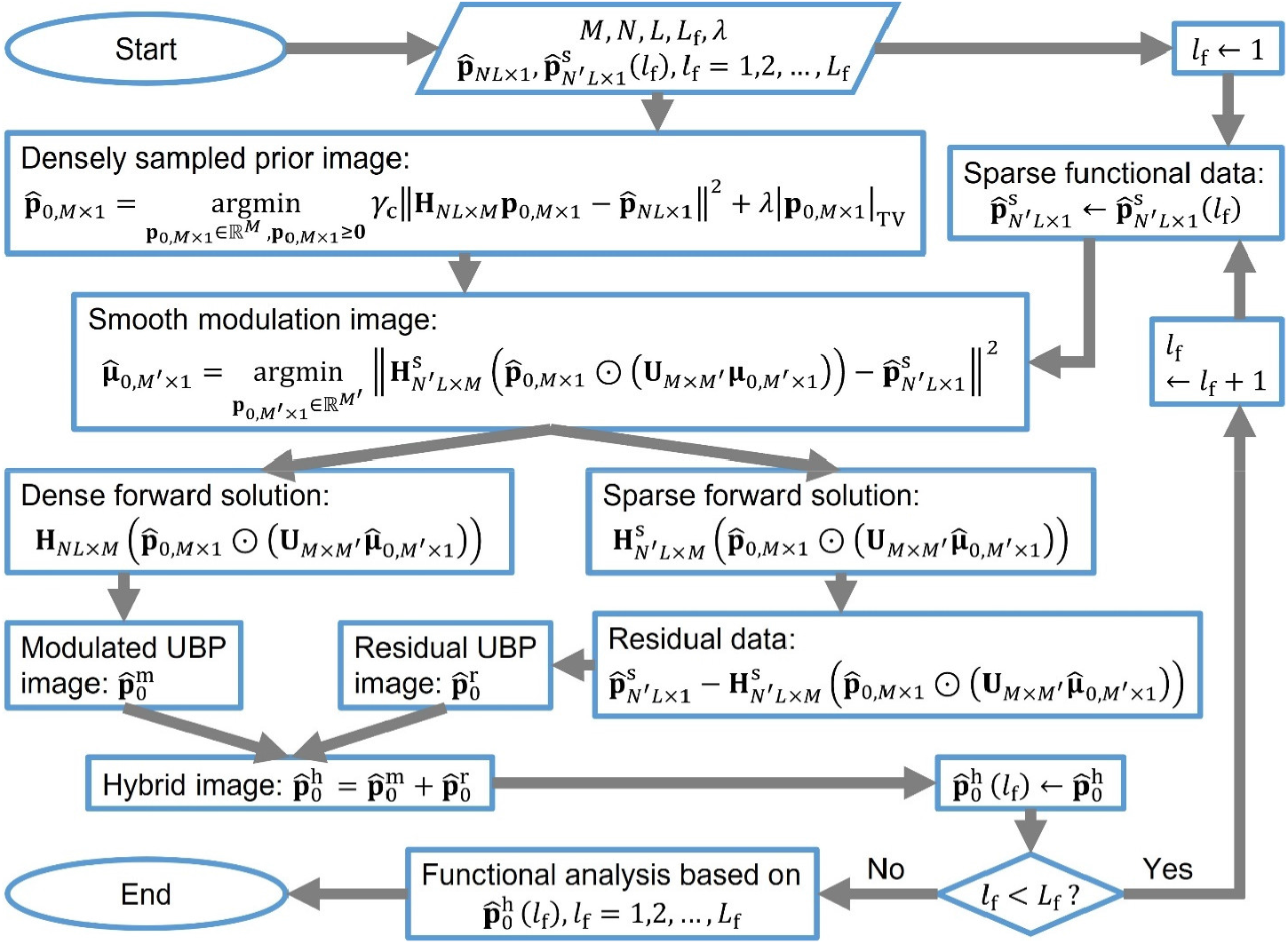
Workflow of the hybrid method for image reconstruction from sparsely sampled raw data and a prior image.

**Supplementary Fig. 7.**
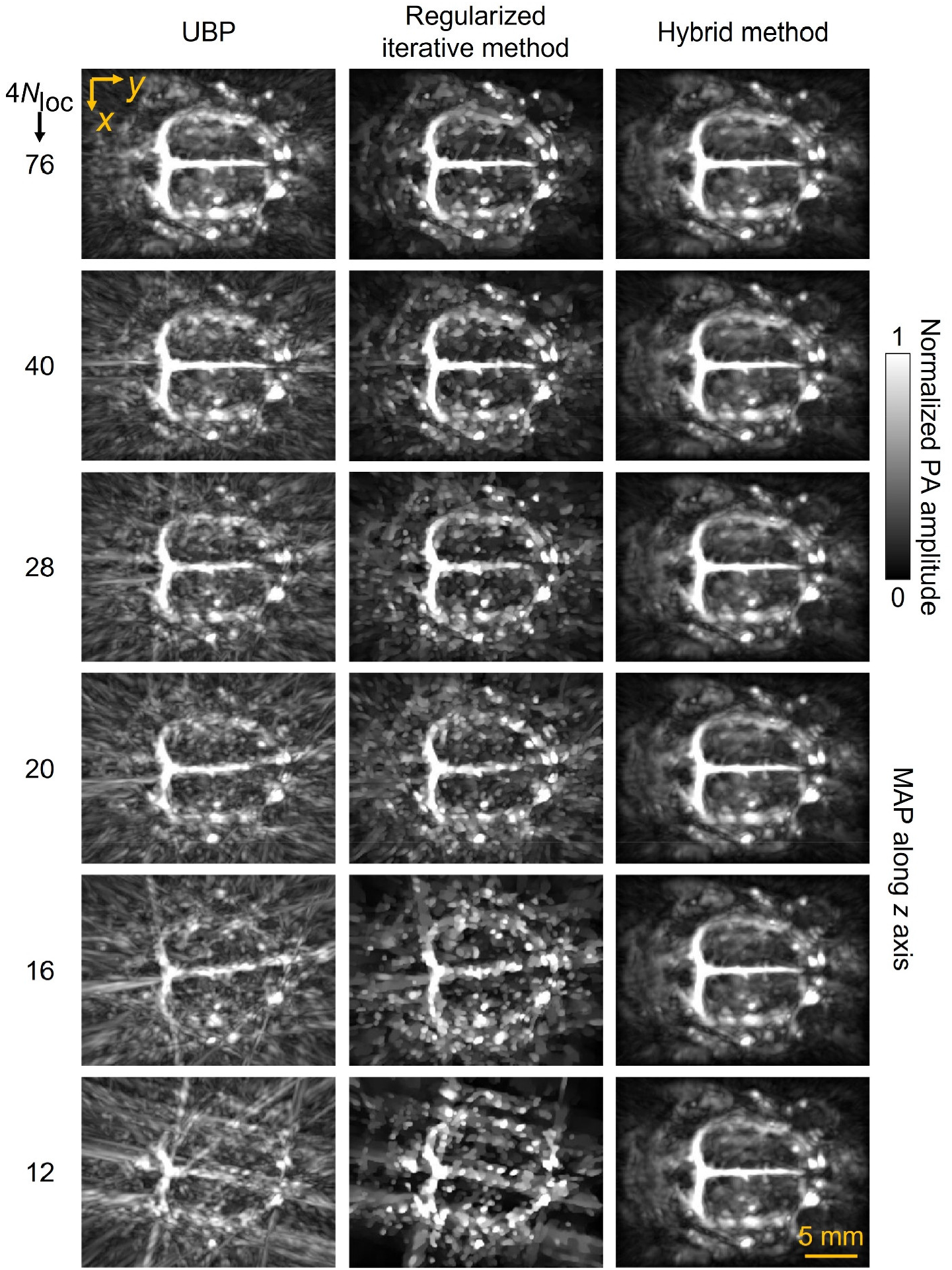
MAPs of the images shown in **Fig. 2a**.

**Supplementary Fig. 8.**
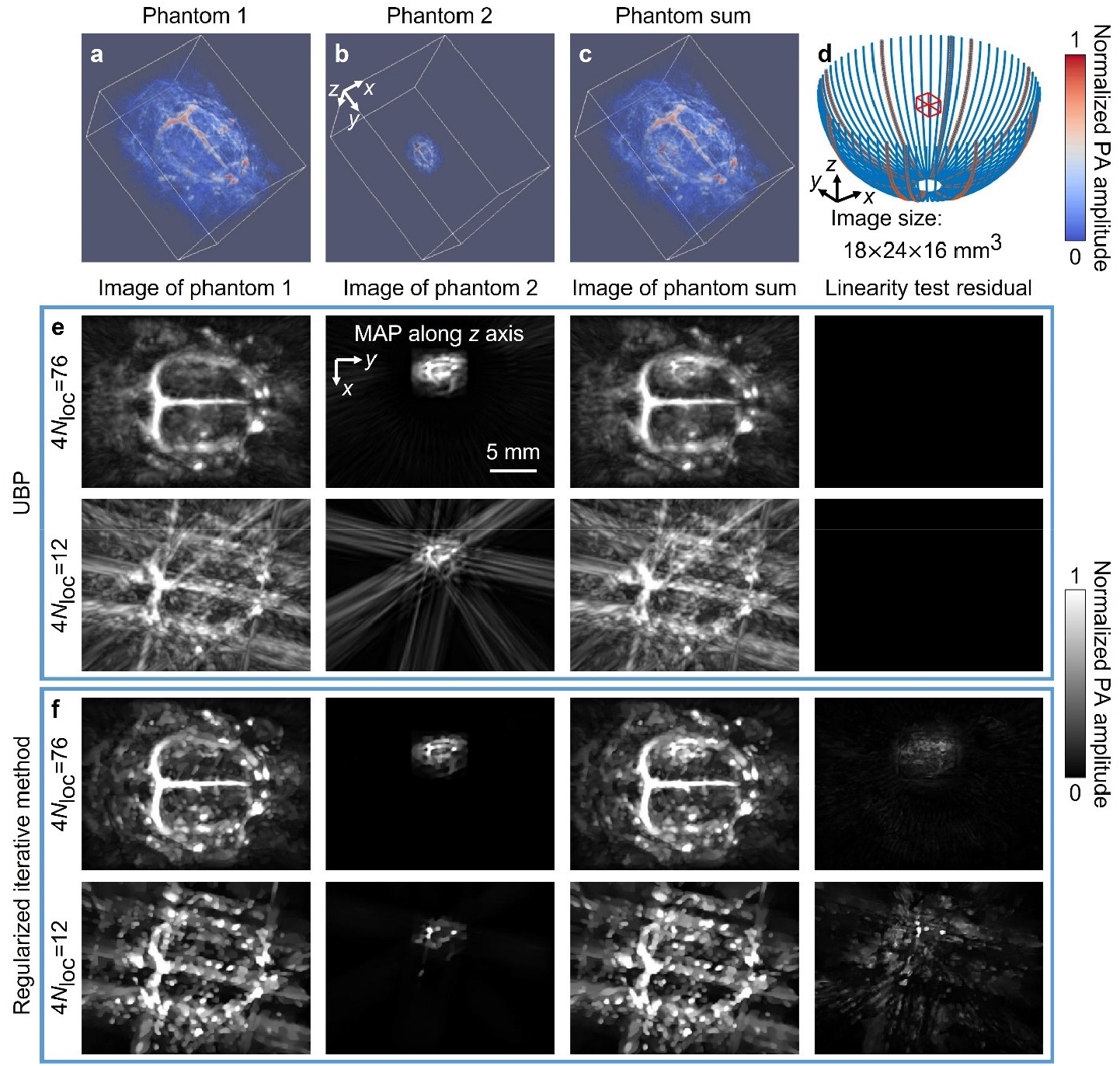
Linearity test of UBP and regularized iterative method. **a–c** The first and second numerical phantoms and their sum, respectively. **d** The image domain (a rectangular cuboid with red-solid edges) and two virtual arrays: all arcs (4*N*_loc_ = 76) and the arcs with red boundaries (4*N*_loc_ = 12). **e-f** MAPs of the images reconstructed using UBP and the regularized iterative method, respectively, from signals detected by the two virtual arrays, and the MAPs of the linear test residuals.

**Supplementary Fig. 9.**
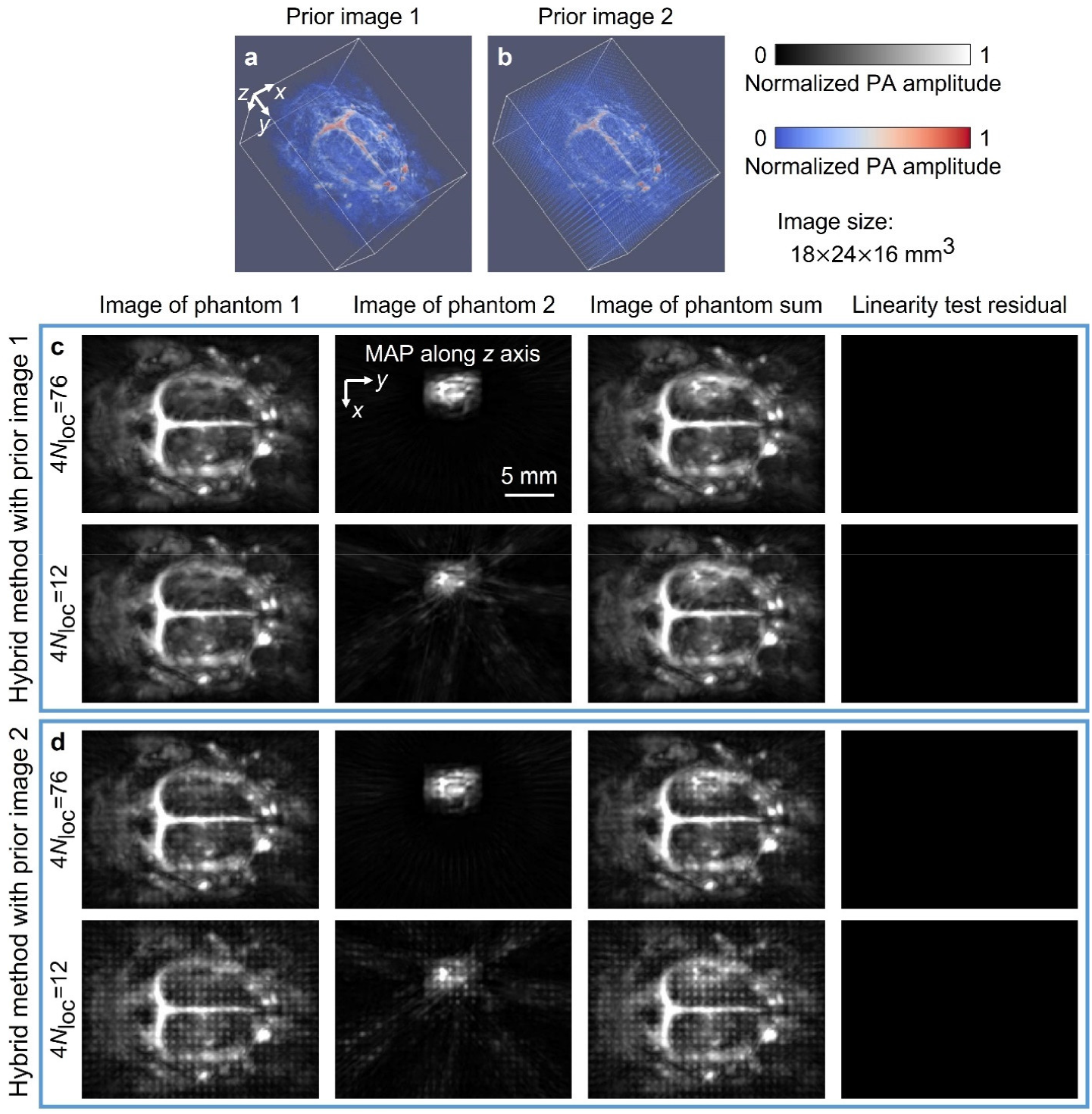
Linear test of the hybrid method with two prior images and the signals used in **Supplementary Fig. 8. a** The ground-truth image used as the first prior image. **b** The second prior image obtained by adding scattered point sources to the first one. **c-d** MAPs of the images reconstructed using the hybrid method with the first and second prior images, respectively, from signals detected by the two virtual arrays (4*N*_loc_ = 76 and 4*N*_loc_ = 12), and the MAPs of the linear test residuals.

### Supplementary Note 7 Numerical simulations of hybrid method for fast functional imaging

**Supplementary Fig. 10.**
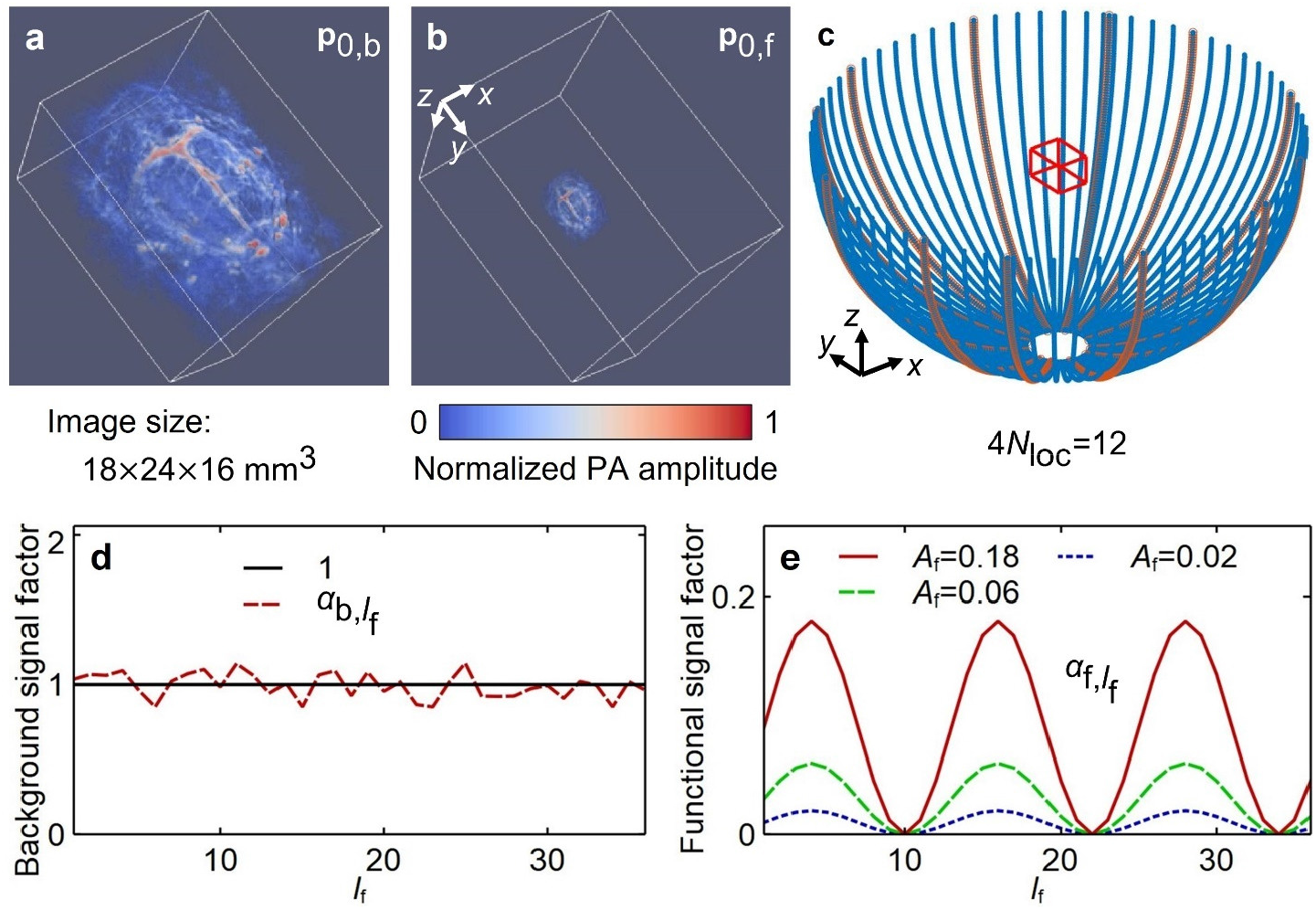
Numerical phantoms for functional imaging. **a-b** The background and functional numerical phantoms, respectively, for functional imaging simulation. **c** A virtual array formed by 12 arc arrays, shown as arcs with red boundaries. **d-e** Modulation factors of the background and functional phantoms, respectively.

We obtain numerical phantoms for functional imaging using

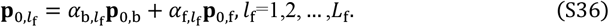

Here, 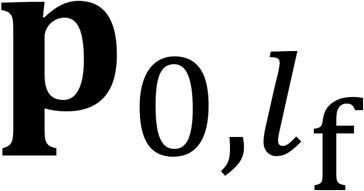 (**Supplementary Fig. 10a**, voxel size 0.1 × 0.1 × 0.1 mm^3^) is the *l*_f_ -th numerical phantom; *p*_0,b_ (**Supplementary Fig. 10b**) is the background phantom obtained in imaging with dense sampling and *p*_0,f_ is the functional phantom obtained from by smoothing, downsampling, and zero padding *p*_0,b_ ; 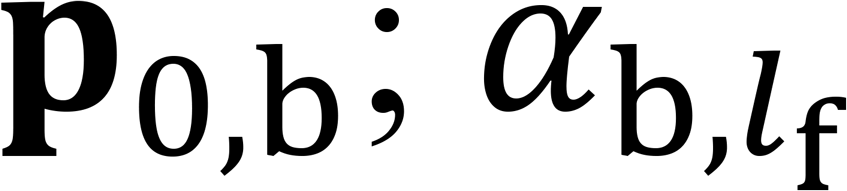 and 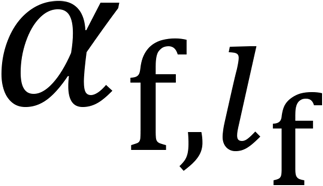 are modulation factors of the two phantoms, respectively; and *L*_f_ is the number of numerical phantoms for functional imaging. The way *p*_0,b_ is obtained guarantees that the mean value of nonzero voxels in *p*_0,_ approximately equals that in *p*_0,f_. For simulations in this study, we use a virtual array formed by 12 arc arrays (**Supplementary Fig. 10c**), let *L*_f_ = 36, and let 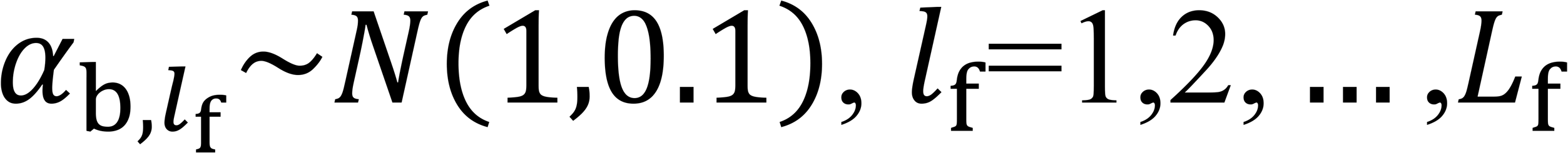 (**Supplementary Fig. 10d**), an amplitude similar to the image relative difference we observed in mouse brain functional imaging. Also, we let

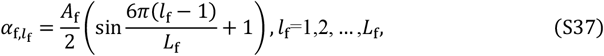

where *A*_f_ is the functional amplitude. The values of 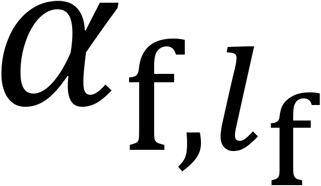 with *A*_f_ = 0.18, 0.06, 0.02 are shown in **Supplementary Fig. 10d** and used in the simulations.

We perform forward simulations, image reconstructions (UBP, the regularized iterative method in Eq. (S34), and the hybrid method in Eq. (5)), and functional signal extractions (Eq. (7)) with different *λ*_f_. The functional images extracted from four sets of images (ground-truth images and images reconstructed with three methods) using the regularized-correlation-based method with *λ*_f_ = 1.6 are shown in **Supplementary Fig. 11** and **Supplementary Video 1**. We observe that the hybrid method is superior to both UBP and the regularized iterative method in functional imaging with sparse sampling. The results with other values of *λ*_f_ also support this observation.

**Supplementary Fig. 11.**
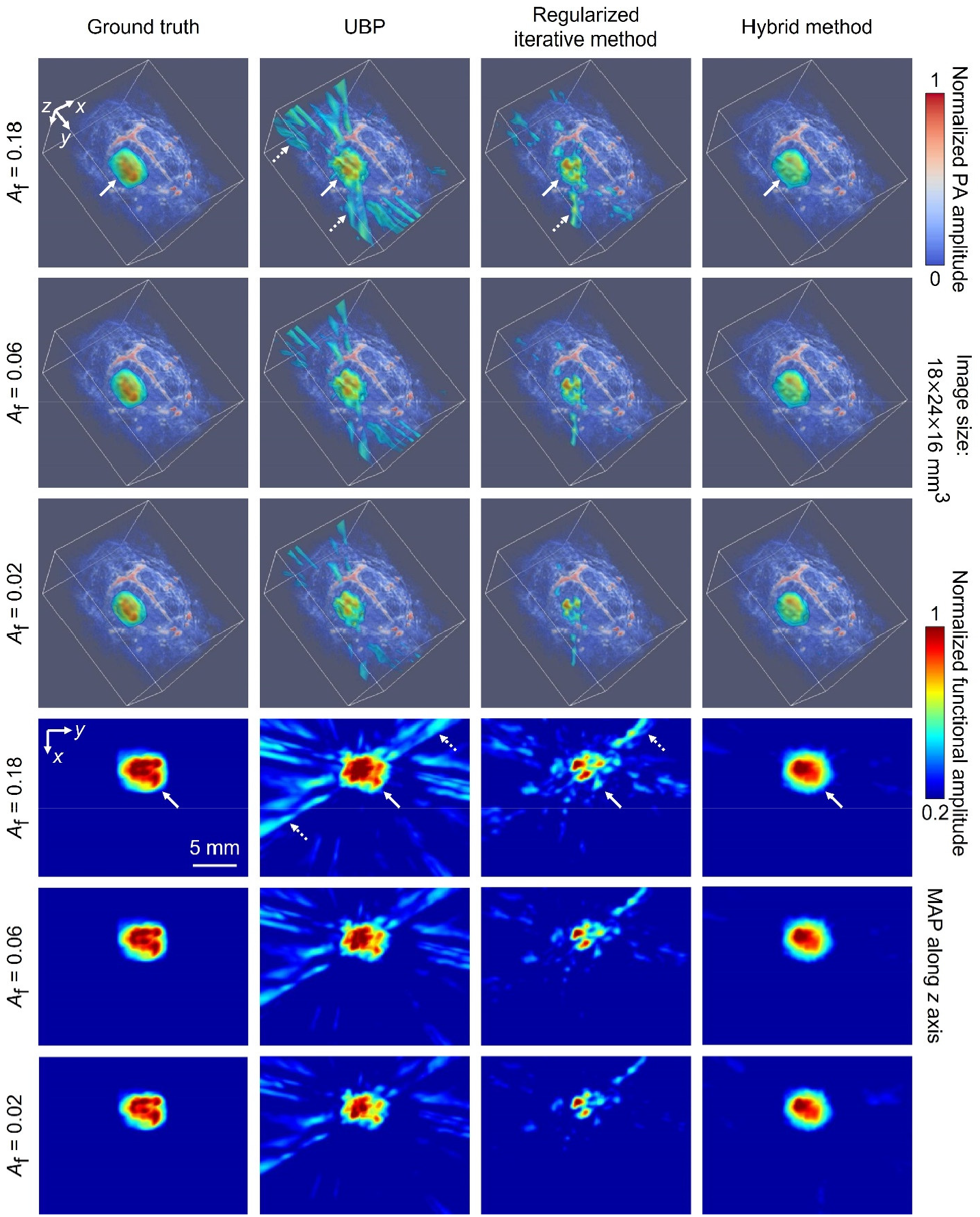
Functional images extracted using the regularized-correlation-based method with *λ*_f_ = 1.6 from ground-truth images and images reconstructed with UBP, the regularized iterative method, and the hybrid method for *A*_f_ = 0.18, 0.06, 0.02. The first three rows show both the 3D functional and background images and the last three rows show the MAPs of the functional images along the *z*-axis. In the first and fourth rows, the true functional regions and examples of false positive regions are indicated by white-solid and white-dotted arrows, respectively.

**Supplementary Fig. 12.**
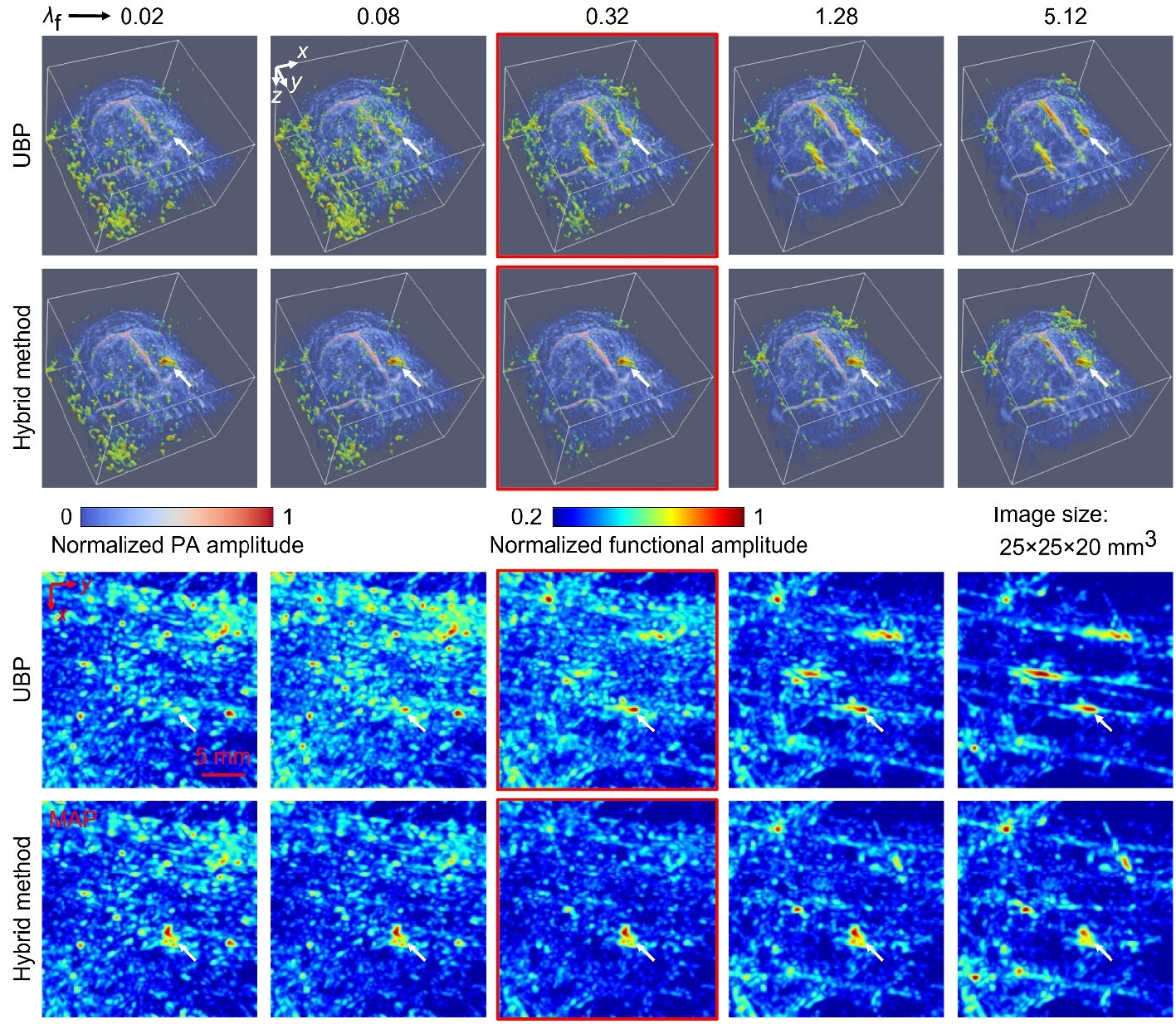
Mouse brain functional images *in vivo* extracted using the regularized-correlation-based method with *λ*_f_ = 0.02, 0.08, 0.32, 1.28, 5.12 from images reconstructed with UBP and the hybrid method (4*N*_loc_ = 12). The first two rows show both the 3D functional and background images and the last two rows show the MAPs of the functional images along the *z*-axis. In all images, the true functional regions are indicated by white-solid arrows, and the other high-amplitude functional regions are false positives. Images for *λ*_f_ = 0.32 are highlighted by red-solid boundaries.

**Supplementary Fig. 13.**
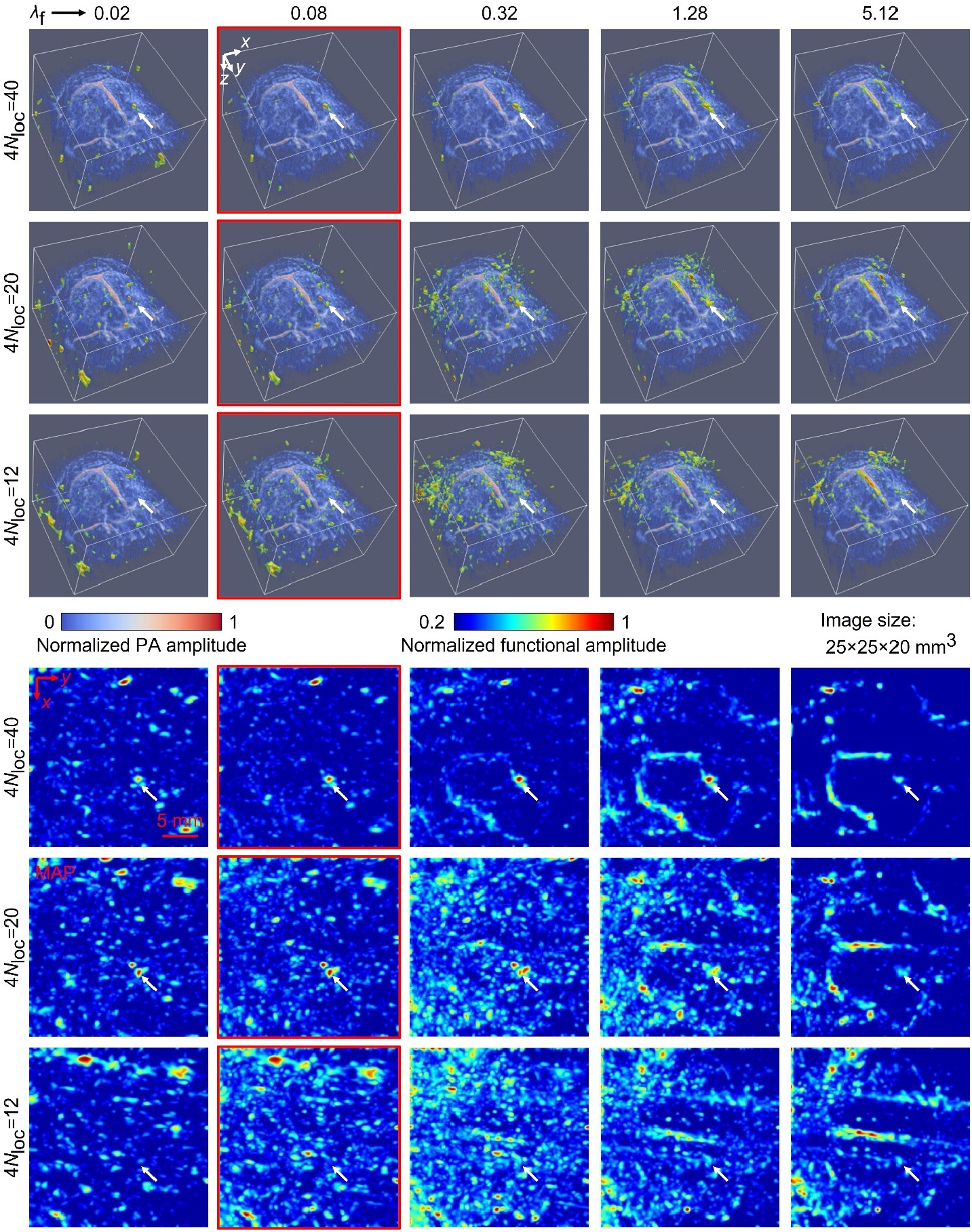
Mouse brain functional images *in vivo* extracted using the regularized-correlation-based method with *λ*_f_ = 0.02, 0.08, 0.32, 1.28, 5.12 from images reconstructed through the regularized iterative method (4*N*_loc_ = 40, 20, 12). The first three rows show both the 3D functional and background images and the last three rows show the MAPs of the functional images along the *z*-axis. In all images, the true functional regions are indicated by white-solid arrows, and the other high-amplitude functional regions are false positives. Images for *λ*_f_ = 0.08 are highlighted by red-solid boundaries.

### Supplementary Note 8 Intra-image nonrigid motion correction

Implementing the system matrix **H** using Eq. (S30) allows for efficient slicing of **H** with respect to image voxel indices and transducer element indices. We propose a method for intra-image nonrigid motion correction through data-driven slicing and manipulation of **H**. In this research, we move a transducer array with *N*_ele_ elements across *N*_loc_ locations to form a virtual 2D transducer array for 3D imaging (with the number of transducer elements *N* = *N*_loc_*N*_ele_). The motion of a source point in the image domain corresponds to temporal shifts of the signals from the source point in the signal domain. To encode the motion information in the forward operator, we modify Eq. (S18) to

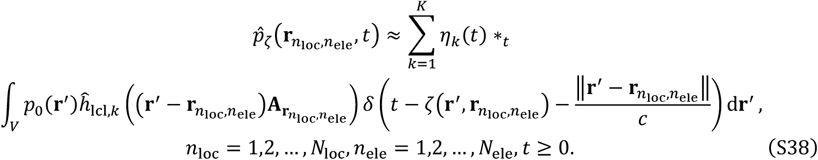

Here, we define 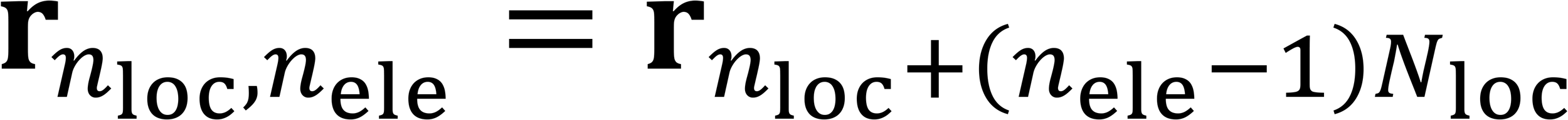 and let 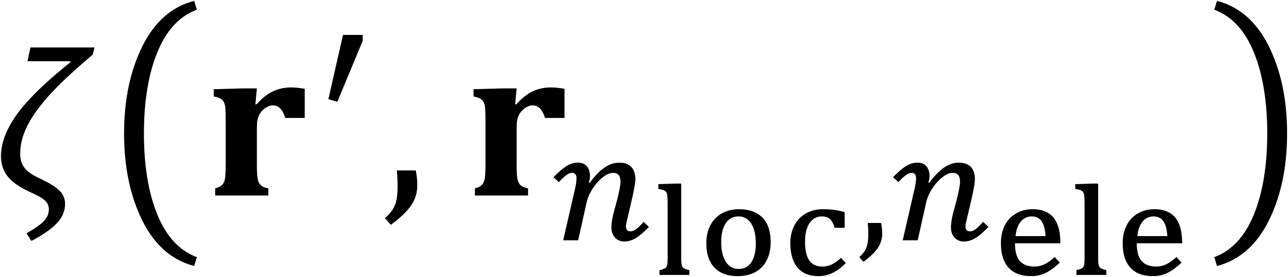 be the motion-induced temporal shift of the signals from the source point at **r**′ that are detected by the *n*_ele_-th element when the transducer array is at the *n*_loc_-th location. It needs to be noted that, due to motion, a source point does not stay at **r**′, and we use **r**′ to represent the source point’s average location.

Next, we mathematically connect the temporal shifts 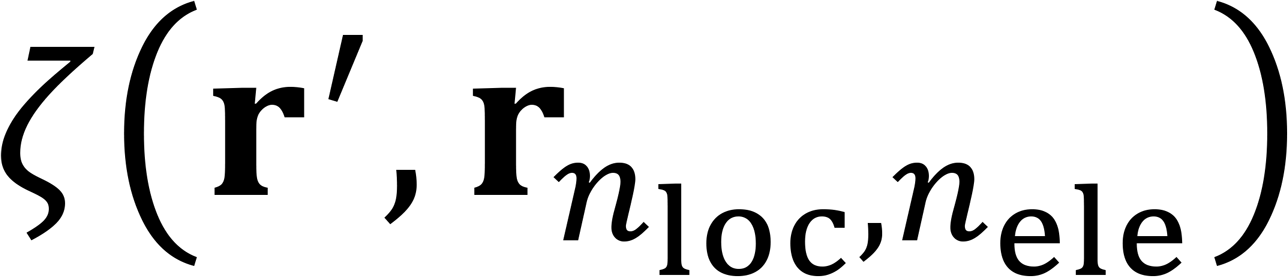 with the motions. For efficiency and robustness, instead of modeling the motions of all source points in the image domain *D*, we model the motions of subdomains 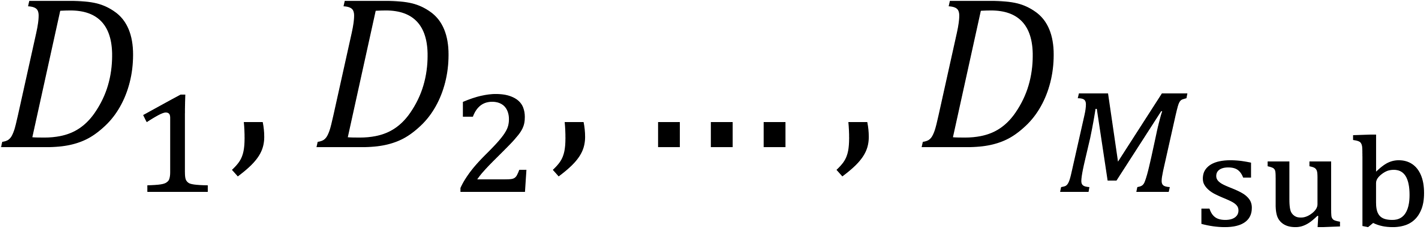 occupying the image domain:

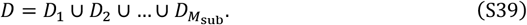

For simplicity, we let all the subdomains be rectangular cuboids of the same size. We denote the center of the *m*_*s*ub_ -th subdomain 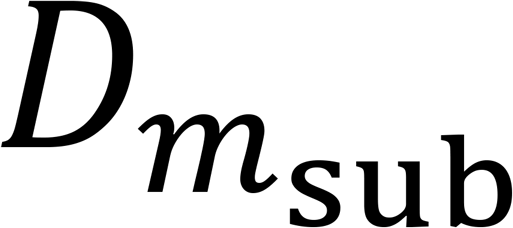 as 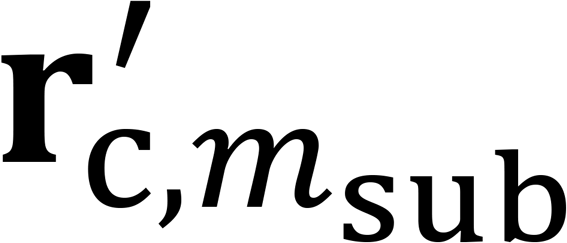 and use the motions of 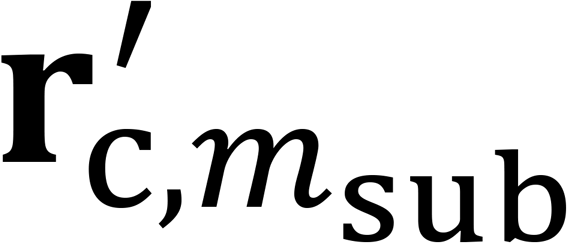 to represent the motions of 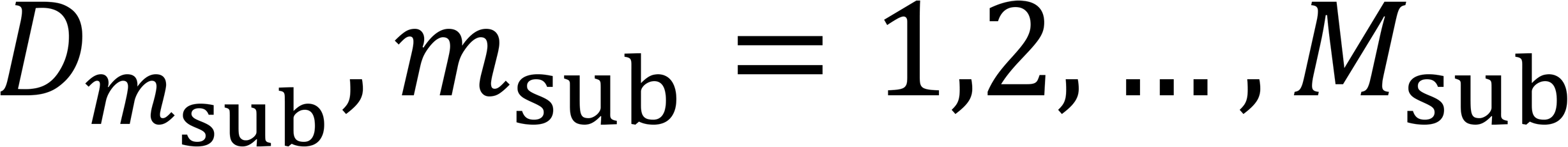. In this research, the motions of the tissues are generally small, and we observe small deformations and rotations of each subdomain. Because of this observation, we ignore the deformations and rotations of each subdomain and only discretize the translations with step sizes *a*_*x*_, *a*_*y*_, and *a*_*z*_ along the *x*-axis, *y*-axis, and *z*-axis, respectively. We represent the motion-induced translation steps of the domain center from 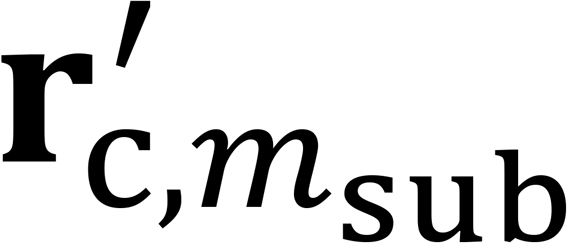 as 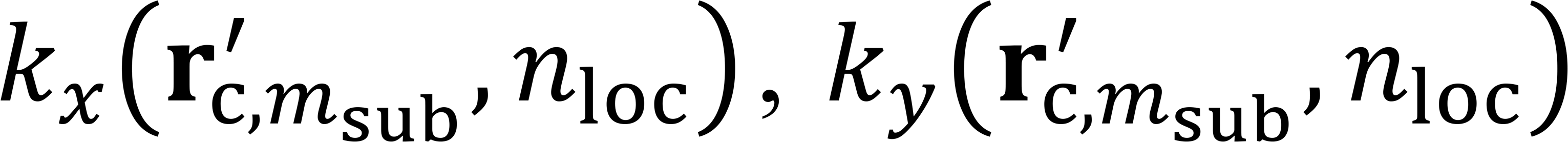, and 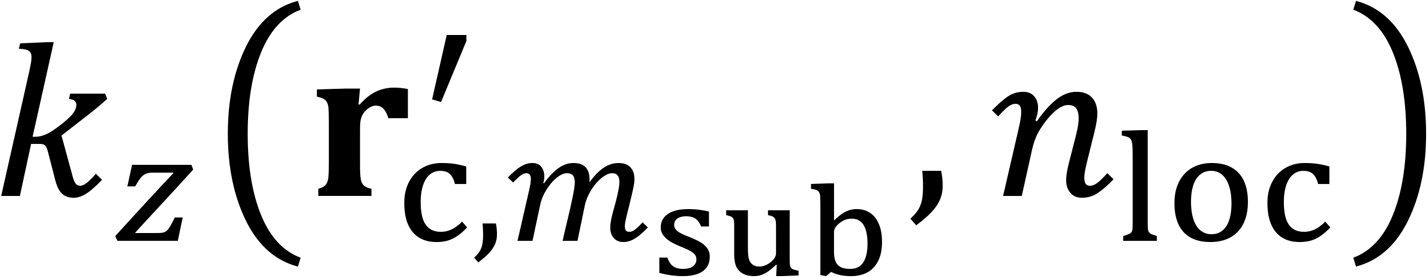 along the *x*-axis, *y*-axis, and *z*-axis, respectively, when the transducer array is at the *n*_loc_-th location. In summary, we express the temporal shifts caused by these translations as

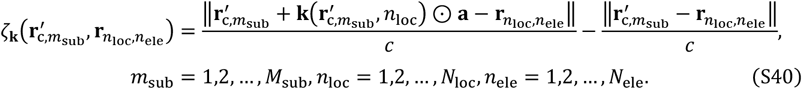

Here, we define a constant vector *a* = (*a*_*x*_, *a*_*y*_, *a*_*z*_) and variable vectors 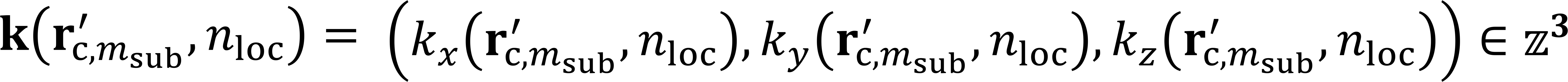, and we group the variable vectors into a set 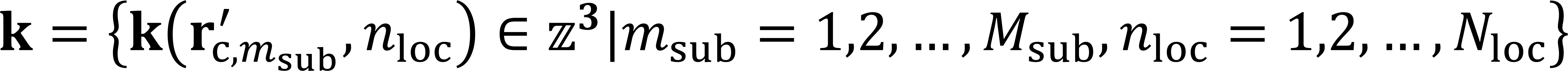. Then we obtain the values of 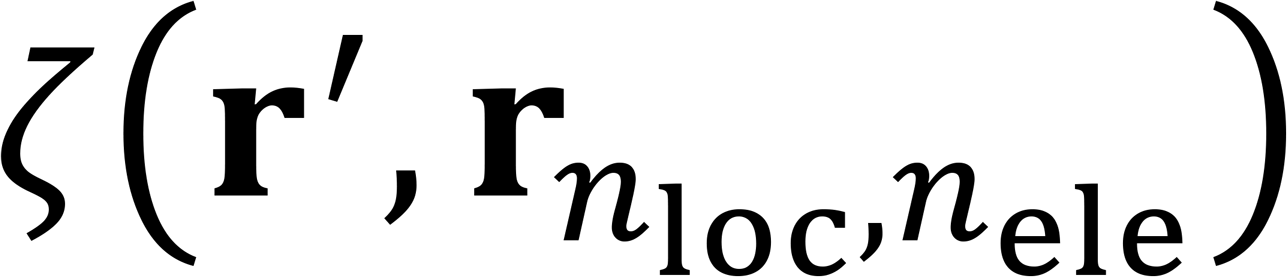 from the values of 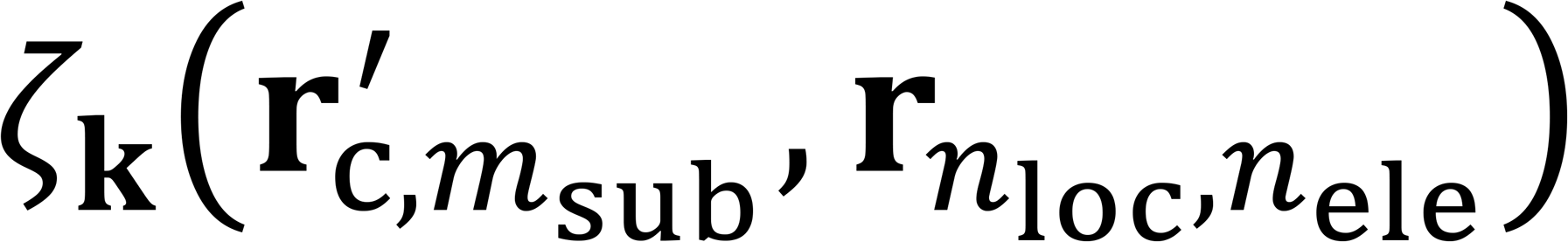 through spatial interpolation in the image domain, which has high accuracy due to the spatial smoothness of the motions of tissues. Thus, we mathematically connect the temporal shifts 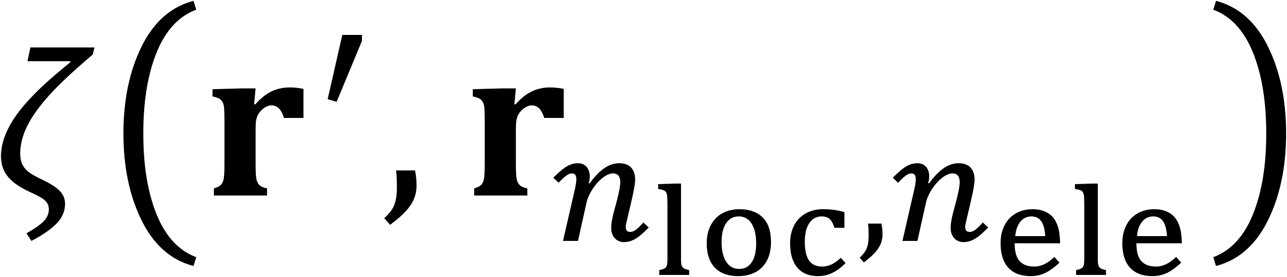 with the motions, described by **k**.

Furthermore, we discretize the motion-incorporated forward operator in Eq. (S38) and propose a motion correction method through Gauss-Seidel-type iterations. We replace 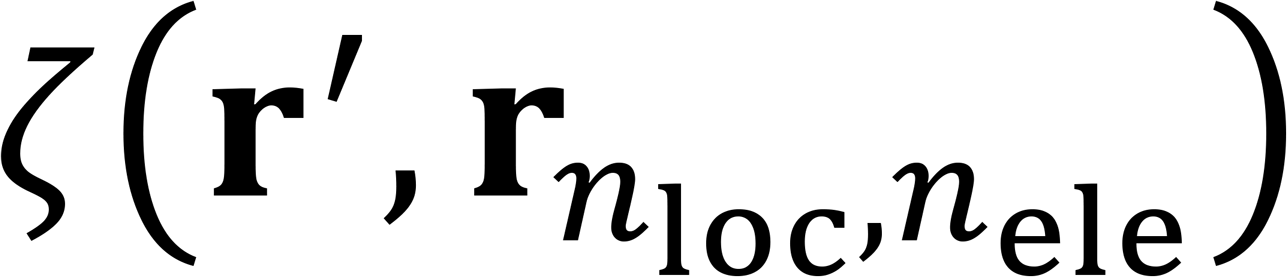 with 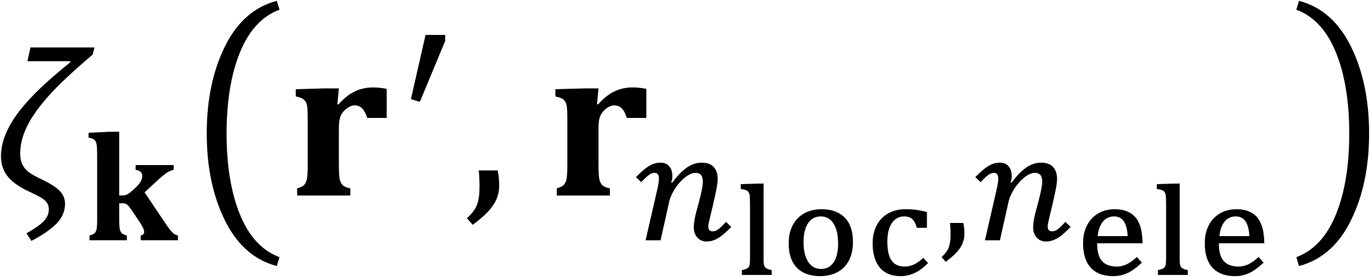 in Eq. (S38) to emphasize these temporal shifts’ dependency on . Then we discretize the forward operator to

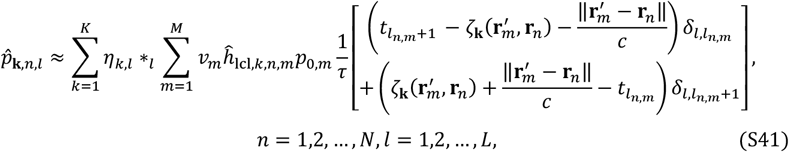

where, *l*_*n,m*_ denotes the temporal index such that 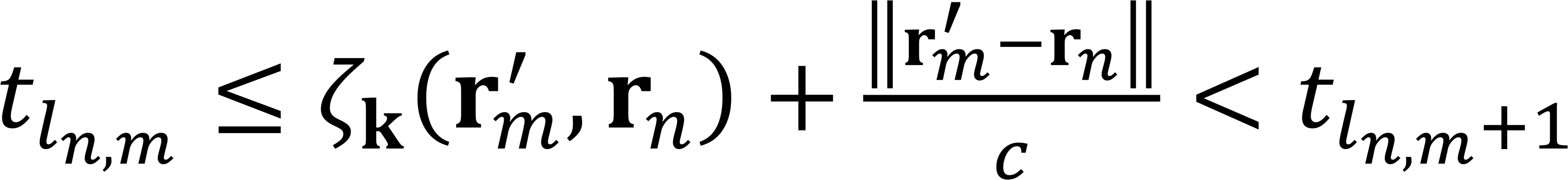. Accordingly, we express the forward operator in matrix form as

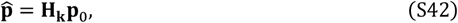

through which we express the data-driven motion correction as a dual-objective optimization problem

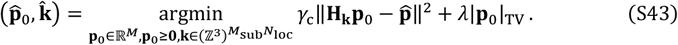

It is challenging to solve this problem directly. Therefore, we simplify it into a convex optimization problem and a combinatorial optimization problem. The convex optimization problem, expressed as

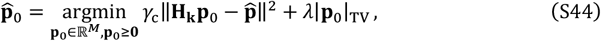

can be solved by the FISTA. The combinatorial optimization problem, expressed as

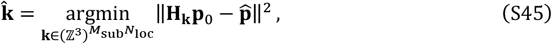

is still challenging.

Instead of solving the problem in Eq. (S45) directly, we solve each vector 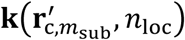 independently without changing other vectors:

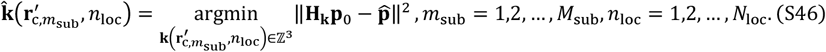

To further simplify Eq. (S46), for each subdomain *D*_*i*_, we decompose *p*_0_ into two images:

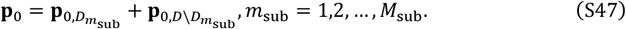

Here, 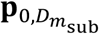 is zero outside the subdomain 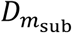 and the same as **p**_0_ inside 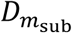, whereas 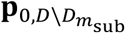 is zero inside the subdomain 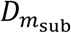 and the same as *p*_0_ outside 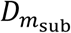. Using this decomposition, we rewrite Eq. (S46) as

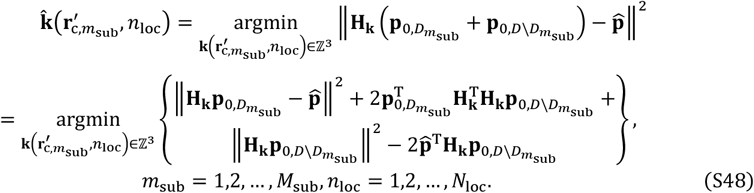

Given the initial pressure 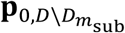, we apply forward simulation (**Hk**) and adjoint reconstruction 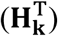 to it to obtain 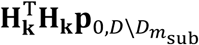, which has small amplitudes inside 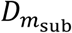 for the detection geometry used in this research. Thus, we have 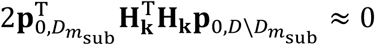. Additionally, although the values of 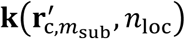 (the motion inside 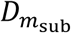) affect 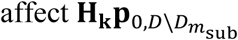 (the signals from the sources outside 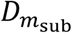) due to spatial interpolation, these effects are minor compared with the values’ effects on 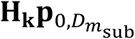. Thus, we ignore 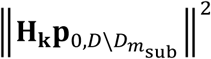 and 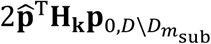 in the optimization for 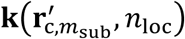. Through these approximations, we simplify Eq. (S48) to

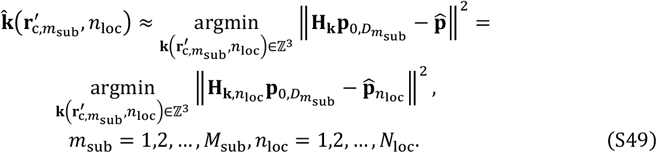

Here, 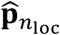 (of shape *N*_ele_*L* × 1) denotes the signals detected by the transducer array when it is at the *n*_loc_-th location whereas signals for other locations are not affected by 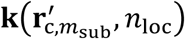, and 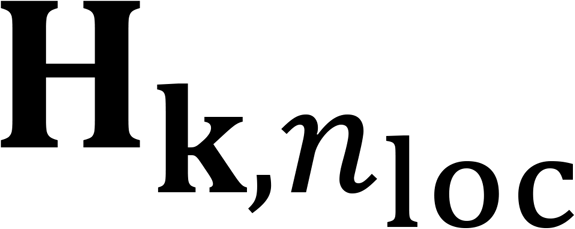 (of shape *N*_ele_*L* × *M*) denotes the corresponding system matrix slices. Additionally, our forward operator allows for forward simulation of any subdomain 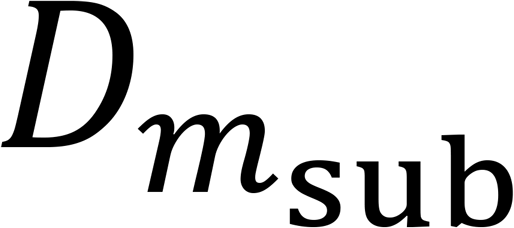 (with *M*_*SD*_ voxels), which is much smaller than the whole image domain *D* (with *M* voxels). Hence, obtaining 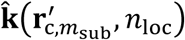 using Eq. (S49) is much more efficient than obtaining it through Eq. (S46). We need to solve the optimization problems in Eqs. (S44) and (S49) in multiple iterations. Thus, instead of obtaining an accurate choice of 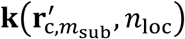 in each iteration, we further decompose the optimization problem in Eq. (S49) into problems

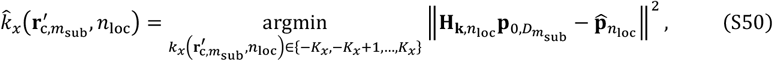

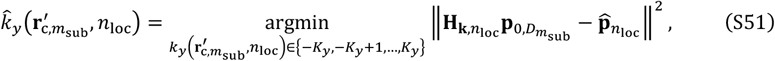

and

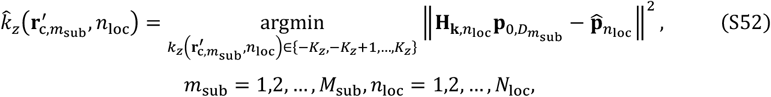

with limited searching ranges determined by parameters *K*_*x*_, *K*_*y*_, and *K*_*z*_, respectively. For each subdomain and each transducer array location, we solve the problems in Eqs. (S50)–(S52) by performing the forward simulation for the subdomain and the transducer array location for 2*K*_*x*_ + 1, 2*K*_*y*_, and 2*K*_*y*_ times, respectively. In summary, the computational complexity of updating all values in **k** once is

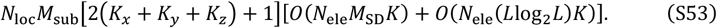

We summarize the workflow of the proposed motion correction method in **Supplementary Fig. 14** and the additional symbols in **Supplementary Table 3**.

**Supplementary Fig. 14.**
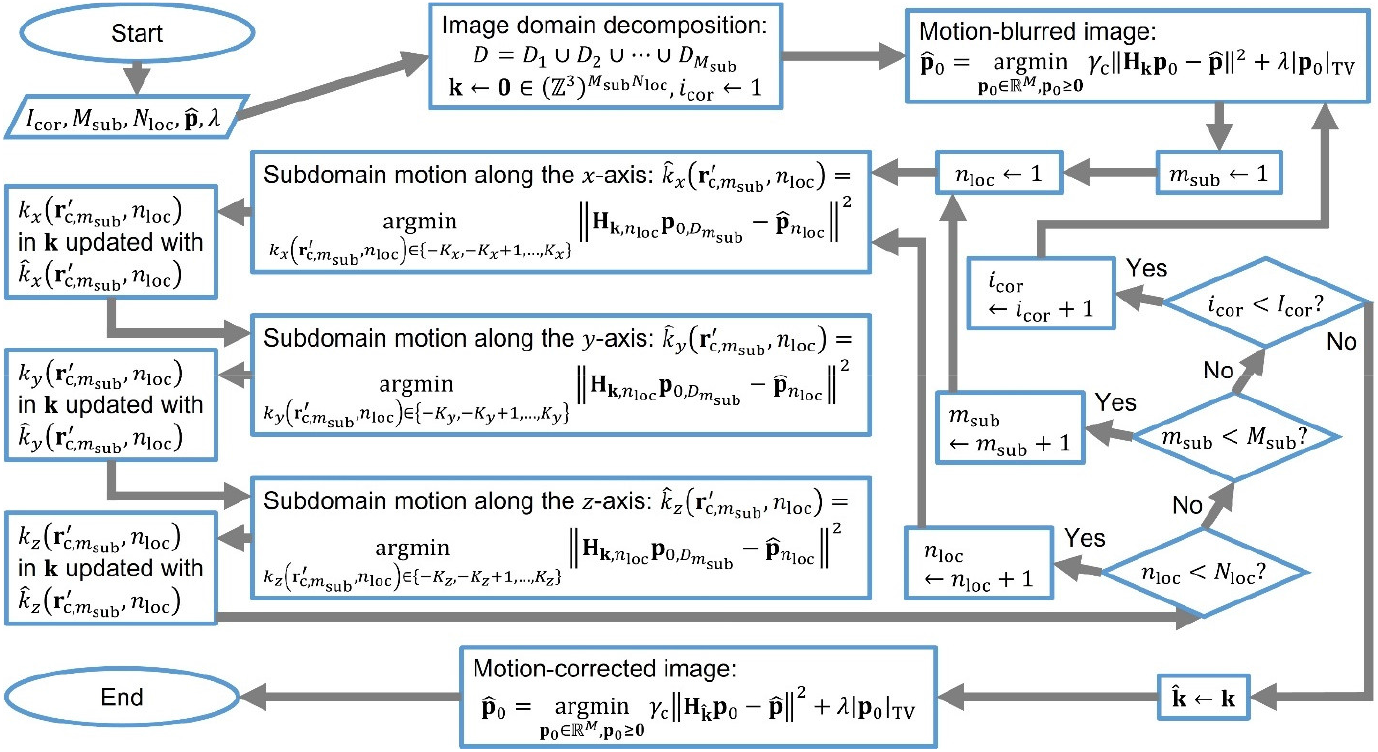
Workflow of the intra-image nonrigid motion correction based on data-driven manipulation of the system matrix.

**Supplementary Fig. 15.**
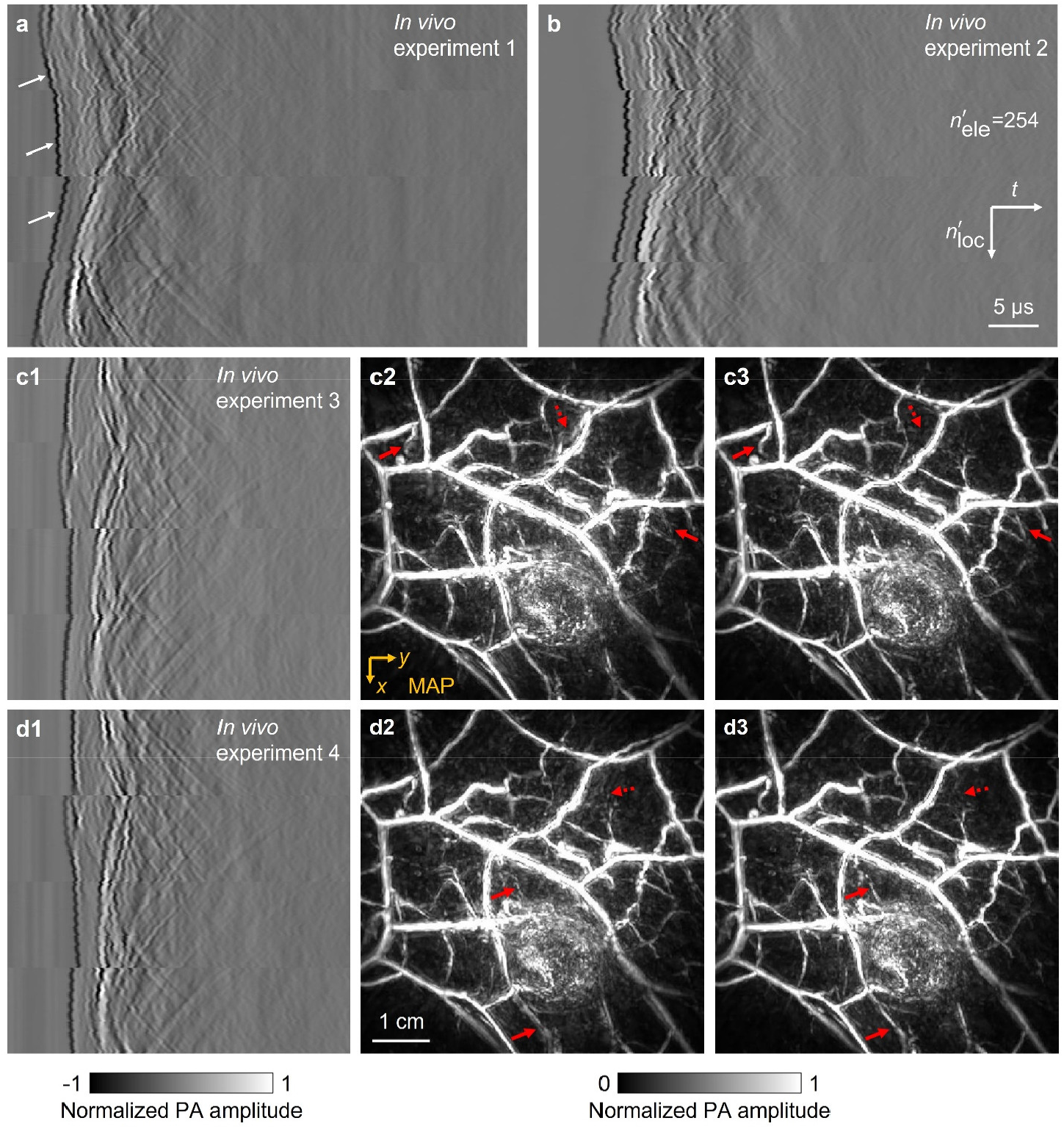
Four sets of signals (*N*_loc_ = 99, *N*_ele_ = 4 × 256) from a human breast *in vivo* with heartbeat-induced motions and reconstructed images. **a-b** Signals in the first two experiments. The reconstructed images are shown in **Fig. 5**. Temporal shifts in **a** caused by the quasiperiodic motion are indicated by white arrows. **c1–c3** and **d1–d3** Signals in the third and fourth experiments, respectively, with the images reconstructed without (**c2-d2**) and with (**c3-d3**) motion correction. Examples of the improvements due to motion correction are indicated by red-solid (feature enhanced) and red-dotted (artifacts suppressed) arrows.

**Supplementary Table 1.**
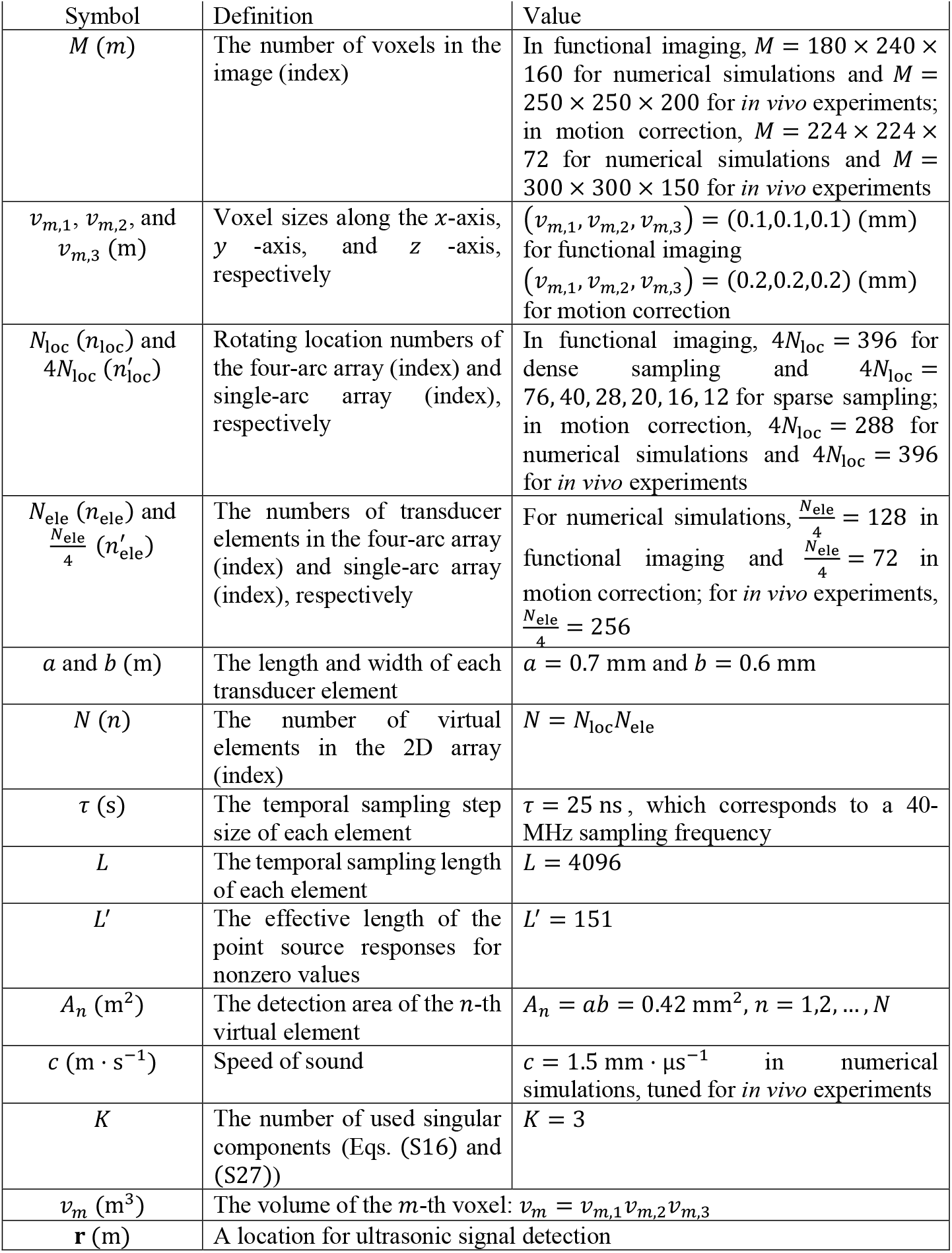
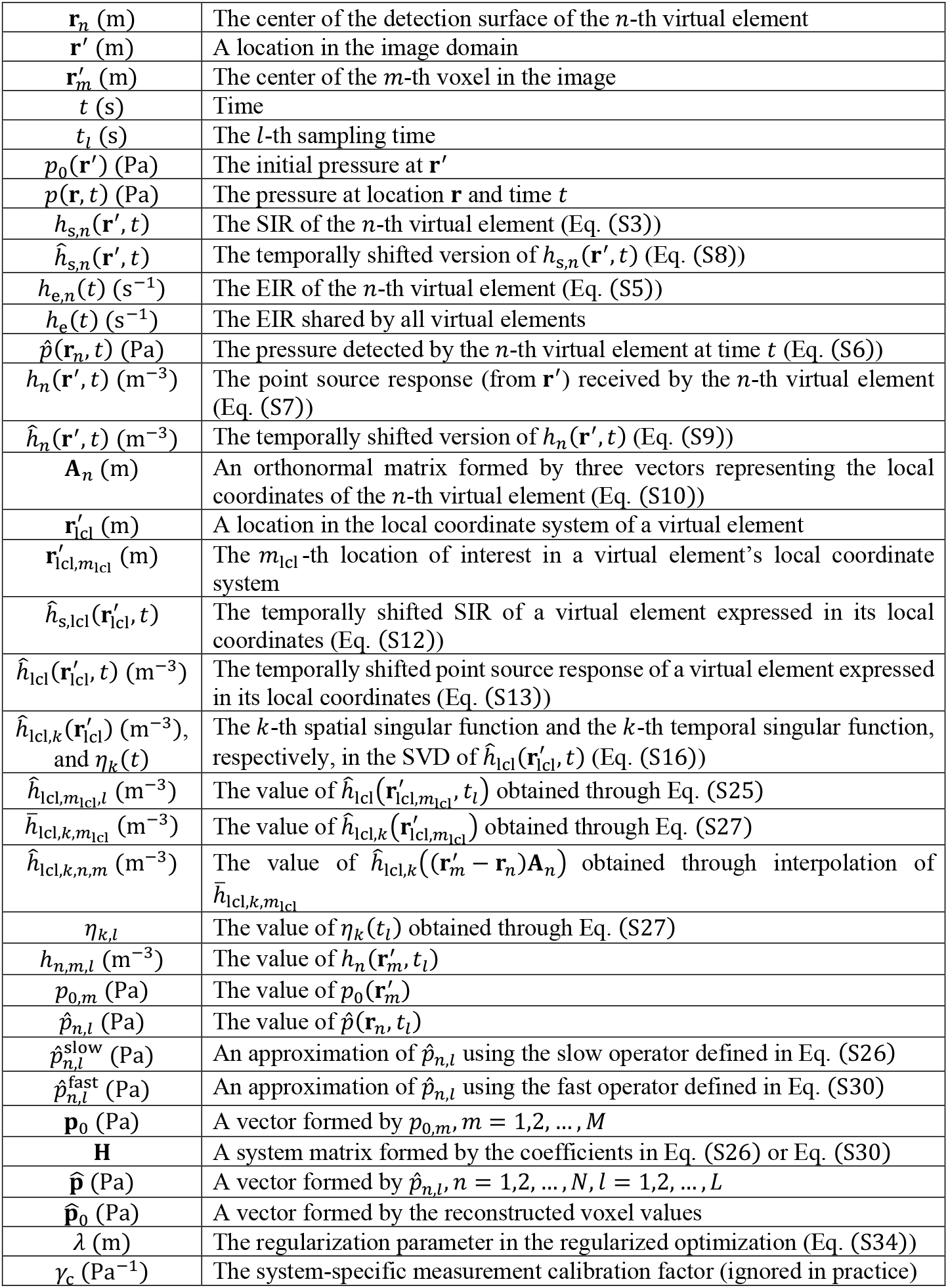
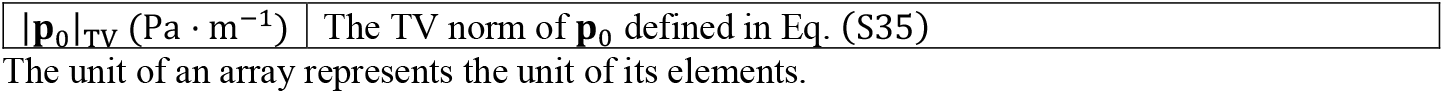
Symbols for the forward operator and image reconstruction.

**Supplementary Table 2.**
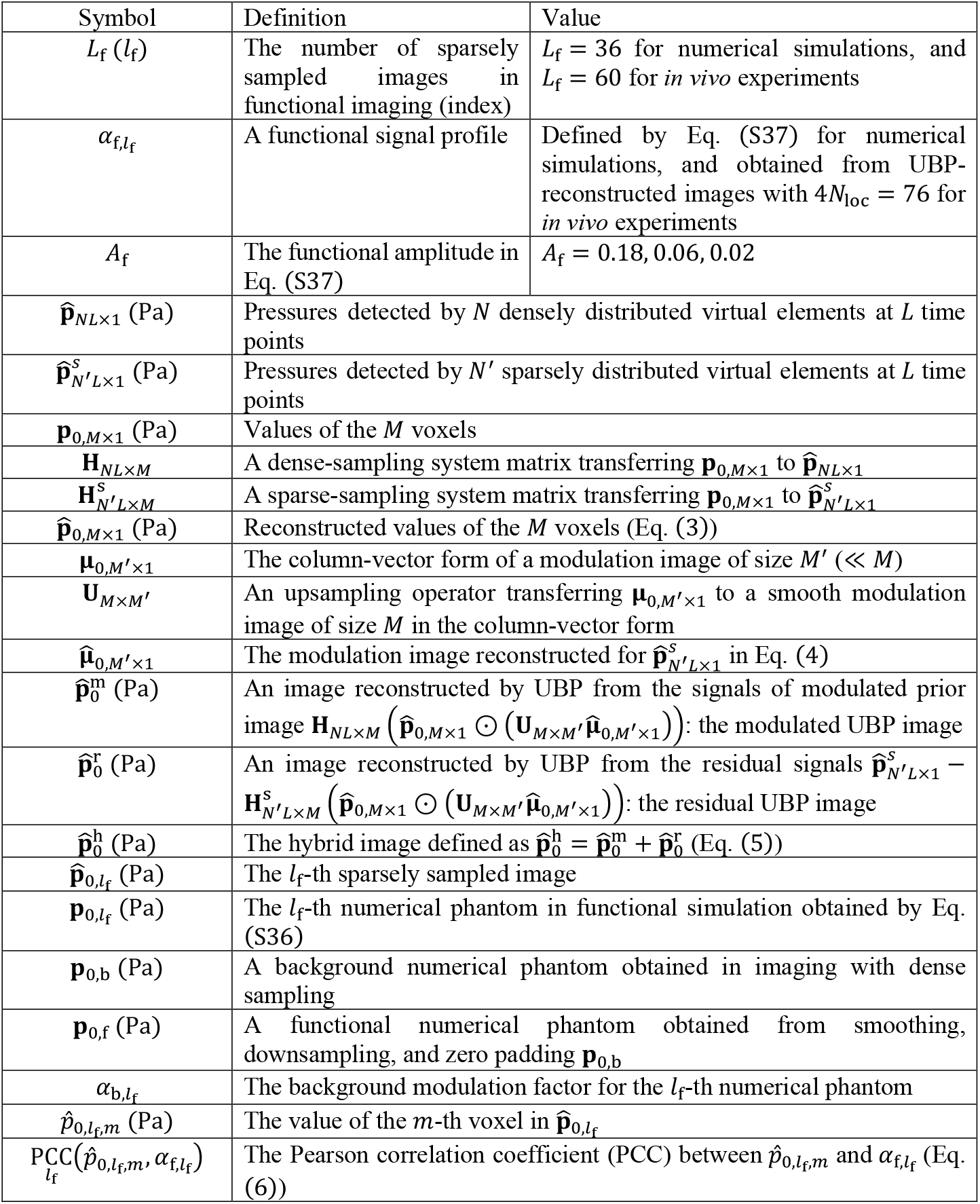
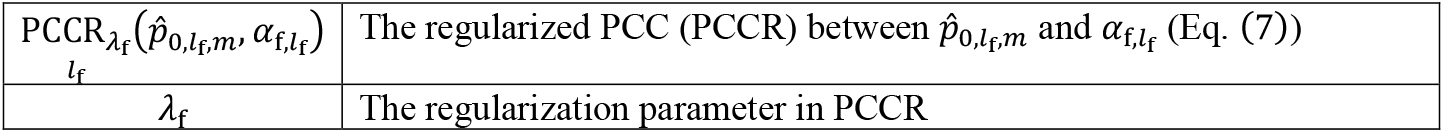
Additional symbols for fast functional imaging with sparse sampling.

**Supplementary Table 3.**
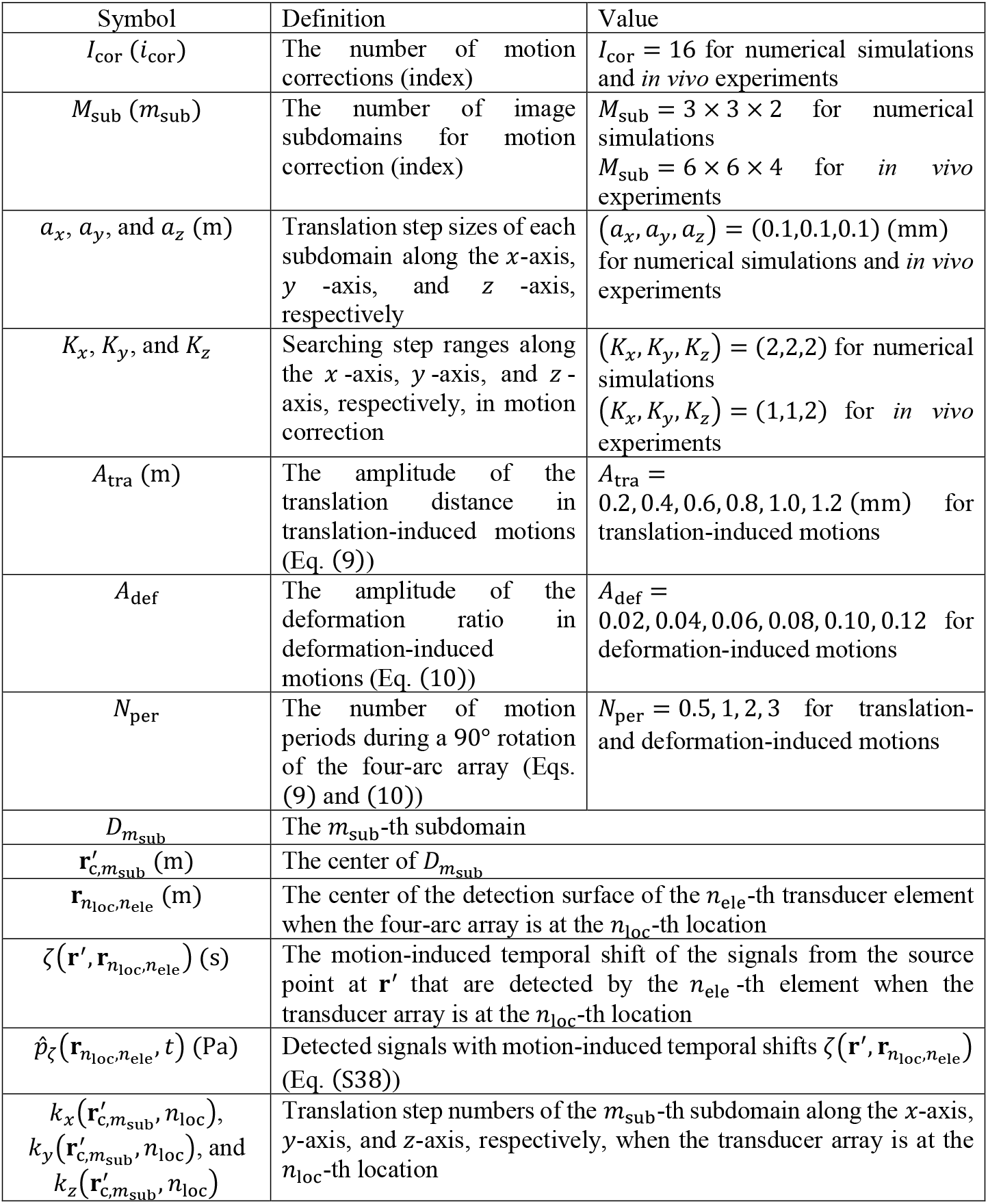
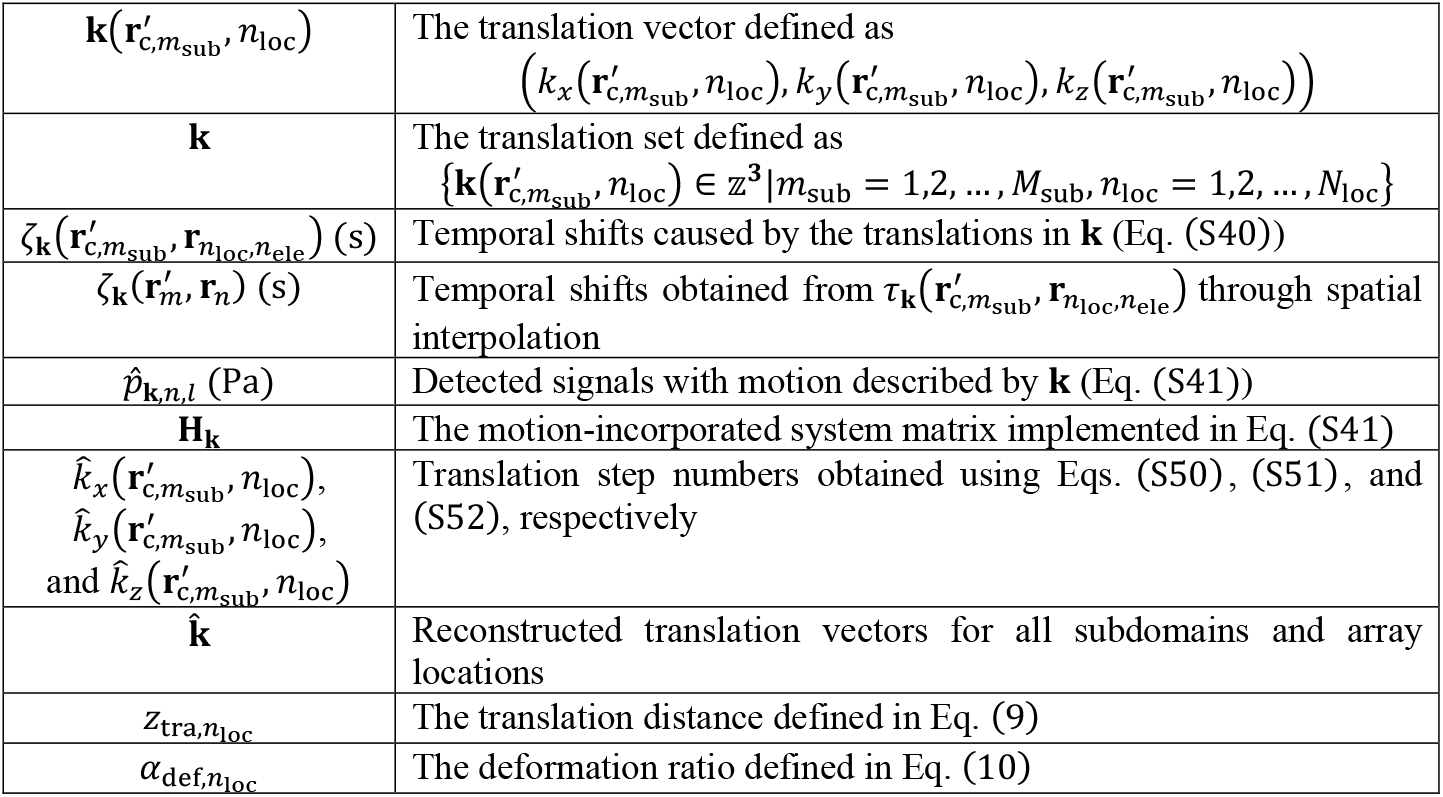
Additional symbols for intra-image nonrigid motion correction.

## Description of Additional Supplementary Files

**Supplementary Video 1** Functional images extracted using the regularized-correlation-based method with *λ*_f_ = 1.6 from ground-truth images and images reconstructed with UBP, the regularized iterative method, and the hybrid method for *A*_f_ = 0.18, 0.06, and 0.02.

**Supplementary Video 2** Mouse brain functional images *in vivo* extracted using the regularized-correlation-based method from images reconstructed with UBP (λ_f_ = 0.32), the regularized iterative method (*λ*_f_ = 0.08), and the hybrid method (*λ*_f_ = 0.32) for 4*N*_loc_ = 40, 20, 12.

**Supplementary Video 3** Synchronized motions of the array and numerical phantom and samples of the detected signals.

**Supplementary Video 4** Translation-induced motions and samples of the detected signals for *A*_tra_ = 1.2 mm, *N*_per_ = 0.5, 1, 2, 3.

**Supplementary Video 5** Deformation-induced motions and samples of the detected signals for *A* _ef_ = 0.12, *N*_per_ = 0.5, 1, 2, 3.

**Supplementary Video 6** MAPs of images reconstructed from signals with translation-induced motions (*N*_per_ = 0.5, 1, 2, 3, *A*_tra_ = 0.2, 0.4, 0.6, 0.8, 1.0, 1.2 (mm)) using the regularized iterative method without and with motion correction. The red-boxed MAPs (*A*_tra_ = 0.6 mm) are shown in **Fig. 4d**.

**Supplementary Video 7** MAPs of images reconstructed from signals with deformation-induced motions (*N*_per_ = 0.5, 1, 2, 3, *A* _def_ = 0.02, 0.04, 0.06, 0.08, 0.10, 0.12) using the regularized iterative method without and with motion correction. The red-boxed MAPs (*A* _def_ = 0.06) are shown in **Fig. 4e**.

**Supplementary Video 8** MAPs (with closed-up subsets) of images reconstructed from signals with translation-induced motions (*N*_per_ = 0.5,3, *A*_tra_ = 0.6 mm) and deformation-induced motions (*N*_per_ = 0.5,3, *A* _ef_ = 0.06) using the regularized iterative method without and with motion correction.

**Supplementary Video 9** MAPs (with closed-up subsets) of human breast images *in vivo* reconstructed using the regularized iterative method without and with motion correction.

